# Nucleotide binding halts diffusion of the eukaryotic replicative helicase during activation

**DOI:** 10.1101/2022.12.23.521684

**Authors:** Daniel Ramírez Montero, Humberto Sánchez, Edo van Veen, Theo van Laar, Belén Solano, John F. X. Diffley, Nynke H. Dekker

**Author notes:** **Materials & Correspondence.** Correspondence and requests for materials should be addressed to N.D. or J.D.

## Abstract

The eukaryotic replicative helicase CMG centrally orchestrates the replisome and leads the way at the front of replication forks^1^. Understanding the motion of CMG on the DNA is therefore key to our understanding of DNA replication. *In vivo*, CMG is assembled and activated through a cell-cycle-regulated mechanism involving 36 polypeptides that has been reconstituted from purified proteins in ensemble biochemical studies^2,3^. Conversely, single-molecule studies of CMG motion have thus far^4–6^ relied on pre-formed CMG assembled through an unknown mechanism upon overexpression of individual constituents^7,8^. Here, we report the first activation at the single-molecule level of CMG fully reconstituted from purified yeast proteins and the quantification of its motion. We observe that CMG can move on DNA in two ways: by unidirectional translocation and by diffusion. We demonstrate that CMG preferentially exhibits unidirectional translocation in the presence of ATP, whereas it preferentially exhibits diffusive motion in the absence of ATP. We also demonstrate that nucleotide binding halts diffusive CMG. Taken together, our findings support a mechanism by which nucleotide binding allows newly assembled CMG to engage with the DNA within its central channel without melting it, halting its diffusion and facilitating the initial DNA melting required to initiate DNA replication.

## Introduction

Eukaryotic DNA replication is catalyzed by a MDa-sized dynamic protein complex known as the replisome. The replisome is powered by the replicative helicase CMG (Cdc45/Mcm2-7/GINS), which centrally orchestrates the other components and leads the way at the front of replication forks^1^. Understanding the motion of CMG on DNA is therefore crucial to our understanding of how cells successfully replicate DNA. *In vivo*, loading and activation of CMG on DNA occur in temporally separated fashion. In *Saccharomyces cerevisiae* in particular, CMG loading occurs at specific sequences known as origins of replication^1^. First, in the G1-phase of the cell cycle, a set of proteins known as ‘loading factors’ scans the DNA until such origins of replication are located, at which inactive single and double Mcm2-7 hexamers are then loaded onto dsDNA^9–12^. In the subsequent S-phase, double Mcm2-7 hexamers are selectively phosphorylated by the cell cycle-regulated Dbf4-dependent kinase (DDK)^13^. Then, a set of proteins known as ‘firing factors’ facilitates the assembly of full CMG by recruiting the helicase-activating factors Cdc45 and GINS to the phosphorylated Mcm2-7 double hexamers^2^. Upon full assembly, CMG must transition from encircling dsDNA to encircling ssDNA, so that it can unwind dsDNA by steric exclusion of the non-translocation strand^14^. This transition is known as CMG activation and consists of two steps. In the first step, ATP binding allows each CMG in a double hexamer to melt 0.6-0.7 turns of dsDNA within its central channel^3^. In the second step, each CMG extrudes one strand of the double helix from its central channel; this final step requires ATP hydrolysis and the action of the firing factor Mcm10^3^. After having extruded one strand of DNA, each activated sister CMG translocates on ssDNA in a 3’-to-5’ direction by hydrolyzing ATP^1,15,16^, allowing the two helicases to bypass and move away from each other^3^, and committing the cell to initiate DNA replication^1^. This entire process requires a minimal set of 36 polypeptides and has been fully reconstituted from purified *Saccharomyces cerevisiae* proteins in ensemble biochemical studies^17^.

To date, the motion of fully reconstituted and activated CMG has been studied in bulk biochemical assays^3,18^ with a temporal resolution of minutes. These ensemble biochemical studies have provided us with important insights into the average behavior of CMG; nonetheless, lower probability behaviors are averaged out in the ensemble readouts. On the other hand, single-molecule studies have the power to isolate and study low probability events with higher temporal resolution^19^; nevertheless, single-molecule studies of CMG motion so far^4–6^ have focused on preformed CMG assembled through a poorly understood mechanism that requires co-overexpression of individual subunits^7,8^. What is more, given that such pre-formed CMG requires an artificial region of ssDNA to bind DNA^6,7^, these studies could not access the intricacies of CMG activation, such as the role of CMG motion during this process. We therefore set out to assemble and activate CMG from purified *Saccharomyces cerevisiae* proteins and study its motion during this process at the single-molecule level.

### A hybrid ensemble and single-molecule assay to visualize fully reconstituted CMG

One of the biggest challenges of studying the motion of fully reconstituted CMG at the single-molecule level is to prevent all the proteins involved from aggregating onto the long (tens of kbp) DNA molecules needed to observe motion of diffraction-limited fluorescent spots^4,6,20^. To overcome this challenge, we developed a hybrid ensemble and single-molecule assay to 1) assemble and activate fully reconstituted fluorescent CMG onto ∼24 kbp DNA molecules containing a natural yeast ARS1 replication origin in an aggregation-free manner; and 2) use a combination of dual optical trapping and confocal scanning microscopy^21^ to image and quantify the motion of fluorescent CMG along DNA molecules held in an optical trap (Fig. 1a-*i*, Extended Data Fig.1a, Methods). To this end, we functionalized both ends of a linear 23.6 kb DNA containing a natural ARS1 origin with digoxigenin and desthiobiotin moieties at both ends. We then bound the functionalized DNA to streptavidin-coated magnetic beads and used it to assemble and activate CMG (Methods). In short, we loaded Mcm2-7 hexamers onto the bead-bound DNA, phosphorylated double Mcm2-7 hexamers with DDK and washed the beads with 300 mM KCl. We then assembled *and* activated CMG for 15 min in the presence of fluorescently labeled Cdc45^LD555^ (Extended Data Fig. 1a), which supports unwinding near WT levels (Extended Data Fig. 1b). Following CMG assembly and activation, we washed the beads again with 300 mM KCl to select for fully mature CMG^2,22^, and ‘paused’ the reaction by removing ATP. DNA:CMG complexes were then eluted from the magnetic beads by competing the desthiobiotin-streptavidin interaction with an excess of free biotin^23^. Following elution, DNA:CMG complexes were tethered between two optically-trapped anti-digoxigenin-coated polystyrene beads, and transferred into a buffer solution containing Mcm10, RPA and either ATP, no nucleotide, or the slowly hydrolyzable ATP analog ATPγS. We then scanned the DNA with a confocal scanning laser and observed fluorescent CMG helicases as diffraction-limited spots on the otherwise unlabeled DNA (Fig. 1a-*ii*). Approximately a third of the trapped DNA molecules contained diffraction-limited fluorescent CMG spots, typically a single one (Fig. 1b). We deduced the number of CMG per diffraction-limited spot by counting the photobleaching steps within each spot (Extended Data Fig. 3, Methods). As this showed that most spots contained 1 CMG (Fig. 1c), it followed that most DNA molecules had a total of 1 CMG (Extended Data Fig. 1c), where *a priori* one might have expected a total closer to 2 or multiples thereof. We cannot attribute our experimentally measured lower number to the labeling efficiency of Cdc45^LD555^, which we measured to be 85 ± 4 %; rather, we attribute it to (a potential combination of) other factors including loss of Cdc45 during the high salt washes and downstream handling, CMG dissociation at nicks^24^ on the DNA during the ensemble activation, or CMG diffusing off the ends of the DNA during elution. Furthermore, it was recently shown that each Mcm2-7 in a double hexamer independently matures into CMG^22^. Thus, we cannot discard the possibility that in our system only one of the two Mcm2-7 hexamers is fully matured into CMG.

**Figure 1.**
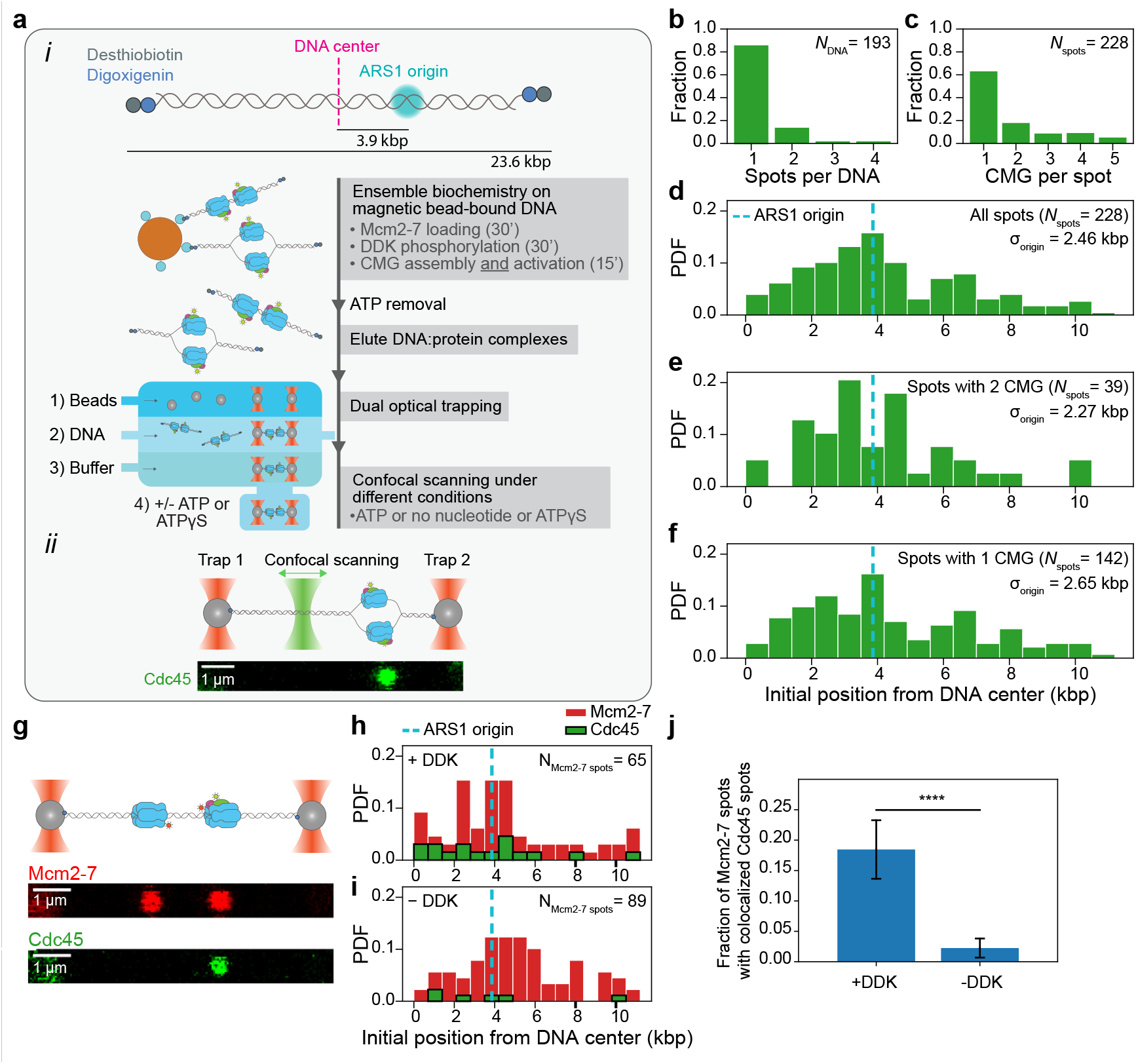
Single-molecule imaging of fully reconstituted CMG helicases. **a-*i***, Description of hybrid ensemble and single-molecule assay to image fully reconstituted CMG helicases. **a-*ii***, Example scan of an optically trapped DNA molecule containing one CMG diffraction-limited spot. **b**, Distribution of numbers of CMG diffraction-limited spots per DNA. **c**, Distribution of numbers of CMG complexes within each diffraction-limited spot. **d-f**, Distribution of initial positions on the DNA of **d**, all CMG diffraction-limited spots, **e**, diffraction-limited spots containing 2 CMG complexes or **f**, diffraction-limited spots containing 1 CMG complex; the ARS1 origin of replication is indicated by the dashed cyan line. **g**, Example scans separately showing Mcm2-7^JF646^ diffraction-limited spots (top) and Cdc45^LD555^ diffraction-limited (bottom) on the same DNA molecule. **h-i**, Distributions of initial positions of Mcm2-7^JF646^ spots and Cdc45^LD555^ spots on the DNA in the **h**, presence or **i**, absence of DDK. In each condition (with and without DDK), the histograms of Mcm2-7^JF646^ <u>and</u> Cdc45^LD555^ initial positions are weighted by the total number of Mcm2-7^JF646^ spots. **j**, Fraction of Mcm2-7^JF646^ diffraction-limited spots with colocalized Cdc45^LD555^ diffraction-limited spots with and without DDK; error bars show the standard error of proportion. Statistical significance was obtained from a two-sided binomial test (p-value=2.2 × 10^−8^).

### Mature CMG is preferentially assembled near origins of replication

We first looked at the initial positions of CMG on the DNA. Of note, because we cannot differentiate between the two possible orientations of the DNA in our experiments, we display the initial positions of CMG in plots showing the distance from the center of the DNA^9^. We observe a wide distribution of initial positions with a peak near or at the ARS1 origin (Fig. 1d). Furthermore, spots containing two CMG complexes are less widely distributed around the origin than spots containing one CMG (Fig. 1e-f). Taken together, these results are consistent with a preferential assembly of sister CMG helicases near the ARS1 origin, followed by the motion of individual activated helicases away from the origin during the 15-min ensemble activation reaction.

### Colocalization of fluorescent Cdc45 and fluorescent Mcm2-7 hexamers is DDK-dependent

Salt-resistant Cdc45 is considered a hallmark of mature CMG^2,22^. Nonetheless, if the Cdc45^LD555^ spots that we observe are part of *bona fide* CMG, their presence on the DNA should be dependent on DDK^2,13^. To confirm this, we quantified the co-localization of red fluorescently labeled Mcm2-7^JF646-Mcm3^ with green fluorescently labeled Cdc45^LD555^ (shown to jointly support DNA unwinding (Extended Data Fig. 2a)) in the presence and absence of DDK (Fig. 1g-j). While nearly 20% of Mcm2-7^JF646-Mcm3^ spots colocalized with Cdc45^LD555^ in the presence of DDK, we observed a ∼4-fold decrease in this colocalization in the absence of DDK (Fig. 1h-j). The 20% colocalization that we observe in the presence of DDK is in agreement with previous observations that the *in vitro* assembly of CMG is less efficient than the loading of Mcm2-7 double hexamers^3,25^. Taken together, these results show that the Cdc45^LD555^ fluorescent spots in our images correspond to *bona fide* CMG. We attribute the residual colocalization of Mcm2-7^JF646-Mcm3^ and Cdc45^LD555^ in the absence of DDK to non-specific interactions (Extended Data Fig. 2h) and/or to traces of phosphorylated Mcm2-7 in the protein preparation^22^.

### Fully reconstituted CMG helicases exhibit two quantitatively distinct motion types

We next sought to quantify the motion of CMG in the presence of ATP. For this, we implemented a change-point algorithm (CPA) to fit linear segments through regions of the position-vs.-time plots of individual spots (Fig. 2a-c, Methods); the slopes of these segments then give us a noise-reduced value of the instantaneous velocities of individual fluorescent spots. To calibrate our analysis, we imaged dCas9^LD555^ with the same imaging conditions that we used for CMG (Fig. 2a, Extended Data Fig. 3); because dCas9^LD555^ is static on the DNA, it provides us with a measure of the velocity error in our system. After drift correction, the distribution of instantaneous velocities of fluorescent dCas9^LD555^ spots after the CPA fit is centered at 0 bp/s and has a width σ_dCas9_ that reflects our experimental uncertainty in velocity measurement (Fig. 2a inset). For all CMG motion analysis, we defined a conservative velocity cutoff of 5 × σ_dCas9_ (= 2.0 bp/s) to categorize fluorescent spots as static or mobile; we considered mobile any fluorescent spot with at least one CPA segment with a slope above this threshold, and all other spots static.

**Figure 2.**
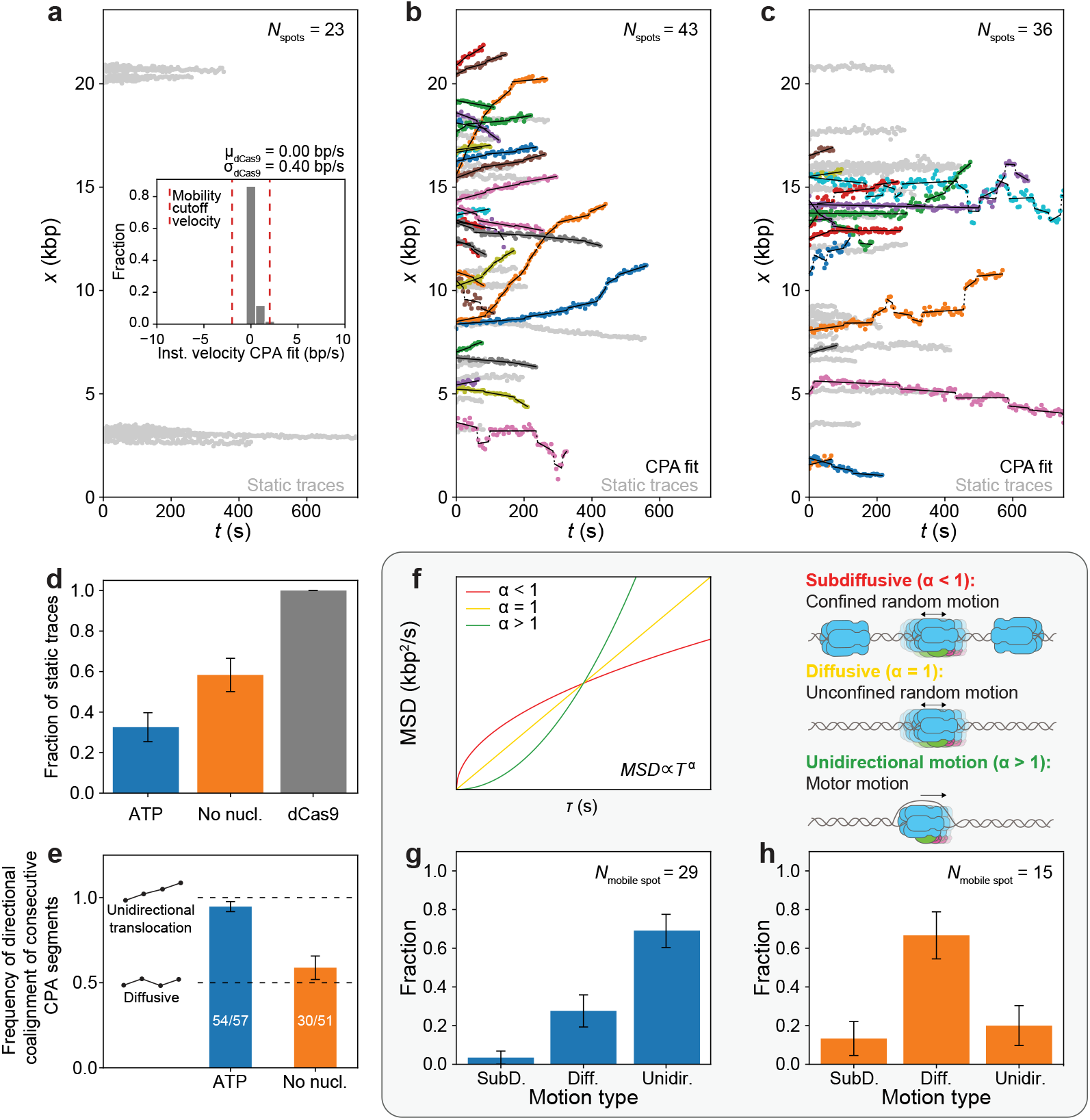
Fully reconstituted CMG exhibits two different motion types. **a**, Position *vs*. time of dCas9^LD555^ spots; (inset) distribution of instantaneous velocities coming from the CPA fits of dCas9^LD555^ spots; red lines show the instantaneous velocity cutoff (5σ_dCas9_) used to separate CMG spots in **b**, and **c**, into static or mobile; CPA fits are not shown for clarity. **b-c**, Position *vs*. time plots of CMG spots in the **b**, presence of ATP **c**, absence of nucleotide; CPA fits are plotted in black, static traces are shown in light gray. **d**, Ratio of static CMG traces in the presence of ATP, absence of nucleotide, and static dCas9 traces. **e**, Frequency of consecutive CPA segments with the same direction for CMG spots in the presence of ATP (blue) or absence of nucleotide (orange); inset diagrams illustrate expected segment directions of a unidirectionally moving spot (top) or a diffusive spot (bottom). **f**, (left panel) Idealized examples of MSD *vs*. delay time τ plots with an anomalous coefficients α < 1 (red), α = 1 (yellow) and α > 1 (green); (right panel) diagrams illustrating the types of CMG motion corresponding to each of these three cases: constrained diffusion (α << 1), free diffusion (α ≈ 1) or unidirectional motion (α >>1). **g-h**, Fraction of mobile CMG traces classified into different motion types in the **g**, presence of ATP or **h**, absence of nucleotide.

Following the approach described above, we determined that ∼70% of CMG spots are mobile when imaged in a buffer solution containing RPA, Mcm10 and ATP (Fig. 2b and 2d, Extended Data Fig. 5a). Unexpectedly, when we imaged CMG in a buffer solution containing RPA, Mcm10 and no ATP, we observed that ∼40% of CMG spots were also mobile (Fig. 2c-d, Extended Data Fig. 5d). Nonetheless, we noticed qualitative differences in motion of CMG in the presence and absence of ATP: while CMG seemed to move unidirectionally in the presence of ATP (Fig. 2b, Supplementary Movie 1), it appeared to move in a more random (e.g. diffusive) manner in the absence of ATP (Fig. 2c, Supplementary Movie 2). To quantitatively characterize these two apparently distinct motion types, we employed two independent approaches. First, we looked at the CPA segments of all the traces in each condition, and calculated the probability that consecutive segments have the same direction (Fig. 2e); in the absence of noise, this probability should equal 1 for unidirectional motion, and 0.5 for random motion. As seen in Fig. 2e, our measured probabilities closely match these expected values, providing quantitative underpinning of our initial observations. As an independent approach, we conducted anomalous diffusion analysis of the mobile traces in each condition (Fig. 2f-h, Extended Data Fig. 5b, 5e and 5h). For each individual trace, we calculated the mean-squared displacement (MSD) as a function of the lag time τ, and then fitted the result to the equation MSD(*τ*) ∝ *τ*^*α*^ to extract the anomalous diffusion coefficient *α* (Methods). The value of *α* then allowed us to classify each trace into different motion types, as *α* ≫ 1 for unidirectionally moving molecules, *α* ≈ 1 for a freely diffusive molecules, and 0 < *α* ≪ 1 for molecules undergoing constrained diffusion (Fig. 2f). This anomalous diffusion analysis confirms that unidirectional motion is most likely when ATP is present (Fig. 2g), whereas diffusive behavior is most likely when ATP is absent (Fig. 2h).

We note that we observed a small population of seemingly diffusive CMG spots in the presence of ATP, and a small population of seemingly unidirectionally moving CMG spots in the absence of ATP (Fig. 2g-h). We hypothesized that these subpopulations might have arisen from misclassification of short traces^26^. To test this hypothesis, we simulated two populations of single-molecule traces of varying lengths within the range of our experimental data: one population solely consisting of unidirectionally moving traces, and the other population solely consisting of freely diffusive traces (Methods). We then carried out the same anomalous diffusion analysis that we did on the experimental CMG data on both simulated data sets. We observed that the distribution of motion types for the simulated unidirectional traces looked very similar to that of the experimental mobile CMG traces in the presence of ATP (Extended Data Fig. 6a and Fig. 2g), whereas the distribution of motion types for the simulated diffusive traces looked very similar to that of the experimental mobile CMG traces in the absence of ATP (Extended Data Fig. 6b and Fig. 2h). Thus, the results of our simulations suggest that, overall, the mobile traces in the presence of ATP represent unidirectional motion, whereas the mobile traces in the absence of ATP represent diffusive motion.

### Analysis of CMG motor motion

Following this identification of two distinct types of CMG mobility, we investigated both in further depth. We first investigated the unidirectional motor motion of CMG, the motion type that powers the replisome. We thus specifically analyzed the velocities of unidirectionally moving CMG spots in the presence of ATP, which yielded a distribution of instantaneous velocities with a peak at ∼5 bp/s (Fig. 3a), consistent with previous single-molecule studies on pre-formed CMG in the presence of RPA^4^. This distribution has a long tail, reaching up to instantaneous velocities of ∼45 bp/s. These higher velocities are low in probability, suggesting that CMG can only achieve these high velocities in short time bursts. Consistent with this, when we calculated the time-averaged velocity of each CMG spot and examined the resulting distribution (Fig. 3a inset), we did not observe such high velocities. The distribution of time-averaged velocities has a peak at ∼5 bp/s, which is consistent with ensemble biochemical studies of CMG motion^18^. Notably, we did not observe any noticeable backtracking of CMG (Fig. 2b), which is consistent with previous studies suggesting that RPA prevents CMG backtracking by keeping the lagging strand template out of the central channel of CMG^4^.

**Figure 3.**
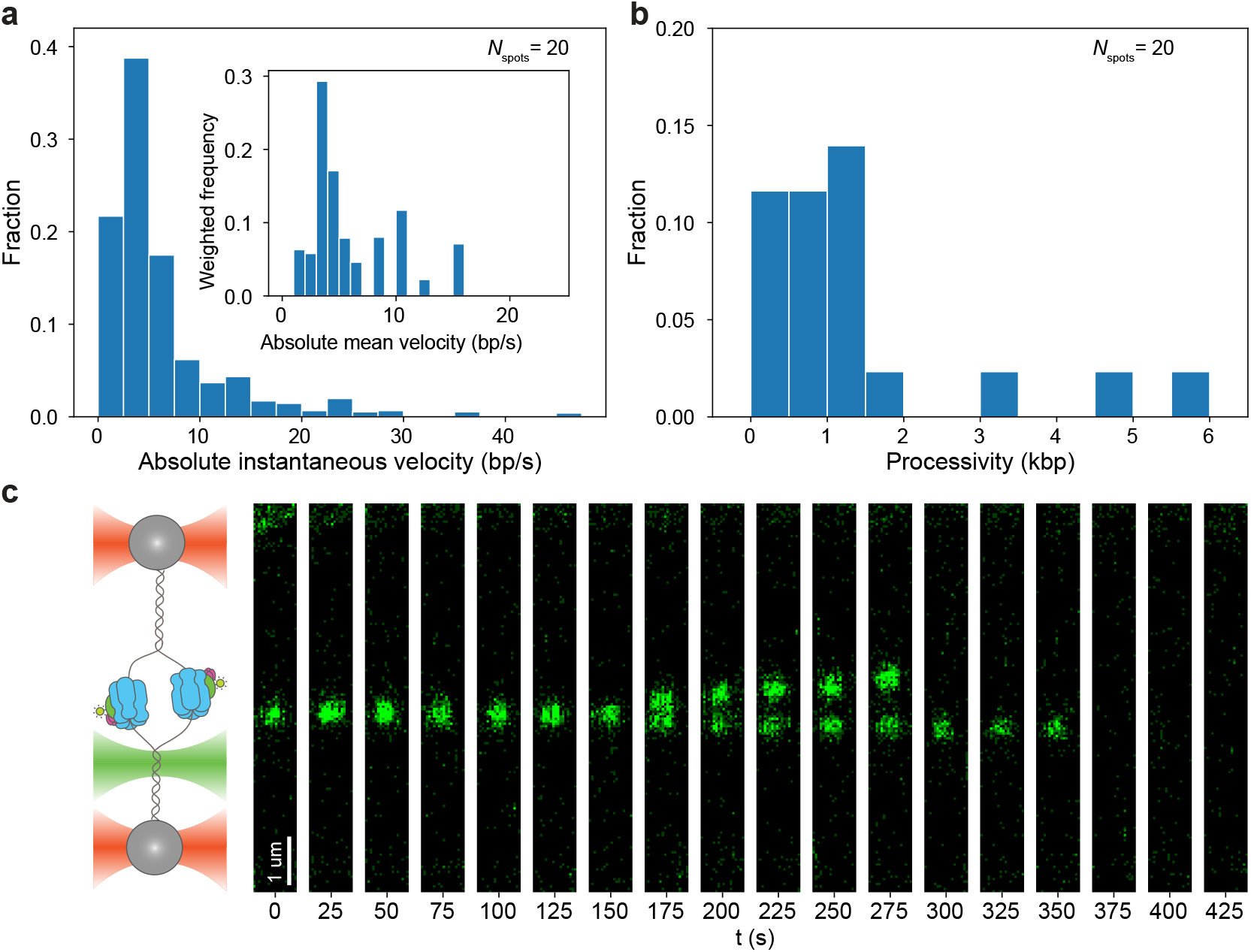
Analysis of CMG motor motion. **a**, Distribution of absolute instantaneous velocities of unidirectionally moving CMG spots in the presence of ATP; (inset) Distribution of absolute mean velocities of unidirectionally moving CMG spots in the presence of ATP normalized by the length of each trace. **b**, Distribution of processivities of unidirectionally moving CMG spot in the presence of ATP. **c**, Example kymograph of two unidirectionally moving CMG spots that start within the same diffraction-limited spot and split up into two distinct diffraction-limited spots that move along the DNA in opposite directions.

When analyzing the unidirectional traces in the presence of ATP, we also observed a few instances of two unidirectionally translocating CMGs initially located within the same diffraction-limited spot, but that then split from one another and give rise to two diffraction-limited spots of half the intensity of the original spot (Fig. 3c, Supplementary Movie 3). These observations are consistent with *in vitro* biochemical studies of helicase activation^3^ showing that sister CMGs move in opposite directions upon their activation. The low probability of these splitting events is to be expected because i) we allow CMG to become activated and translocate on the DNA for 15 min before imaging, and ii) because most DNA molecules contain 1 CMG (Extended Data Fig. 1c).

### Nucleotide binding halts diffusive CMG

Our data shows that CMG diffuses on DNA in the absence but not in the presence of ATP (Fig. 2b-c and 2g-h), suggesting that ATP is involved in stopping this diffusive motion of CMG. To investigate whether it was the binding or the hydrolysis of ATP that stopped the diffusive motion of CMG, we investigated CMG motion in a buffer solution supplemented with RPA, Mcm10 and the slowly hydrolysable ATP analog ATPγS. When we imaged CMG under these conditions, the vast majority CMG spots were found to be static (Fig. 4a-b, Extended Data Fig. 5i). Taken together with our data in the presence and absence of ATP (Fig. 2b-d), our results show that it is the nucleotide binding and not the hydrolysis that halts the diffusive motion of CMG. Furthermore, our results confirm that ATP hydrolysis is required for the unidirectional translocation of CMG^16^.

**Figure 4.**
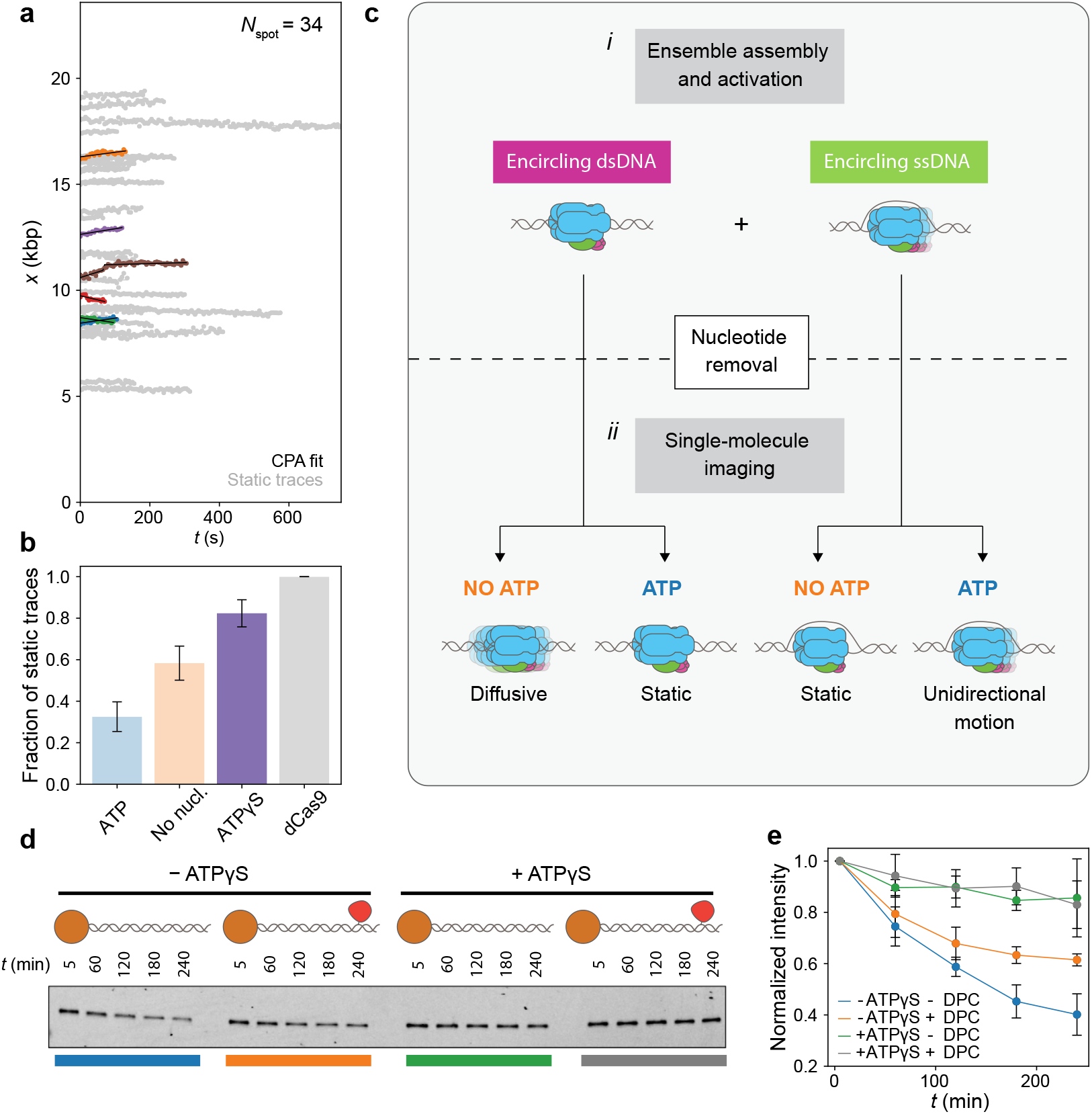
Nucleotide binding halts CMG diffusion. **a**, Position *vs*. time plots of CMG spots in the presence of ATPγS; CPA fits are plotted in black, static traces are shown in light gray and mobile traces are shown in all other colors. **b**, Fraction of static CMG spots in the presence of ATPγS (purple); the results from Fig. 2d, are shown as light bars for comparison. **c**, model proposed to explain experimental motion results; **c**-***i***, proposed two populations of CMG present in the ensemble CMG activation reaction; melted and extruded states correspond to the first and second steps in CMG activation, respectively; **c-*ii***, summary of experimental outcomes in Fig. 2 and Fig. 4a, and proposed explanation of their origins. **d**, Fluorescent scan of an SDS-PAGE gel showing the amount of Cdc45^LD555^ left on linear DNA bound to magnetic beads at one end and containing either a free end or an end capped with a covalently crosslinked methyltransferase. **e**, Densitometry quantification of the experiment shown in **d**, showing the average normalized intensity of three replicates together with their standard deviation. Data points are connected by solid lines to guide the eye.

Previous biochemical studies showed that ATP binding allows newly formed CMG to melt 0.6-0.7 turns of the DNA within its central channel^3^, which was recently confirmed by cryo electron microscopy^25^. Comparing these previous observations with our single-molecule results led us to hypothesize that 1) the diffusive motion that we observed in the absence of ATP corresponded to CMG surrounding dsDNA, and that 2) the halting of such diffusive motion in the presence of ATPγS is due to CMG melting the DNA within its central channel.

To test whether CMG can diffuse onto dsDNA in the absence of ATP, we developed an ensemble CMG sliding assay (Fig. 4d-e, Methods). Briefly, we synthesized two 1.4 kb linear DNA constructs biotinylated at one end and containing an ARS1 origin. The non-biotinylated end of the constructs was then either left as a free end or covalently crosslinked to a M.HpaII methyltransferase^27^. Because the crosslinked methyltransferase is too large to fit inside the central channel of CMG^28,29^, it should stop CMG from diffusing off the end of the DNA, which would otherwise be free to diffuse off the free end of the DNA. We then bound both DNA constructs to streptavidin-coated magnetic beads, and assembled CMG in the presence of fluorescent Cdc45^LD555^ onto them; importantly, we omit the firing factor Mcm10 from the activation reaction to prevent strand extrusion from the central channel of CMG and ensure that CMG is surrounding dsDNA^3^. After CMG assembly, we incubated the bead-bound DNA in a buffer solution with or without ATPγS, and monitored the amount of fluorescent Cdc45^LD555^ present on the DNA over time. As seen in Fig. 4d-e, we detected the fastest decay of Cdc45^LD555^ signal in the DNA construct with a free end in the absence of ATPγS, whereas there was only a small decay in the same DNA construct when ATPγS was present. We also observed some decay, albeit smaller in magnitude, in the DNA construct with a capped end in the absence of ATPγS (Fig. 4d-e); this smaller decay may result from spontaneous opening of the Mcm2-7 ring of CMG in the absence of nucleotide^6,30^ on such timescales. Thus, the results of our ensemble assay support our hypothesis that CMG diffuses on dsDNA in the absence of ATP, and that nucleotide binding halts this diffusive motion.

To test whether the nucleotide-binding-mediated halting of CMG diffusion was due to DNA melting within the central channel of CMG, we employed a recently reported Mcm2 mutant in which six residues directly involved in DNA melting are substituted with alanine (hereafter referred to as Mcm2^6A^)^25^. Notably, Mcm2^6A^ supports CMG assembly but does not support either DNA melting or strand extrusion^25^, allowing us to separate the effect of DNA melting from DNA binding. Thus, if DNA melting by CMG is indeed what halts its diffusive motion, then CMG assembled with Mcm2^6A^ should be fully diffusive in the presence of ATP. Nevertheless, the vast majority of CMG spots assembled with Mcm2^6A^ are static in the presence of ATP (Extended Data Fig. 9a). Furthermore, when we performed our ensemble CMG sliding assay with Mcm2^6A^, we observed the same trends as with WT Mcm2 (Extended Data Fig. 9g-h), providing further evidence that CMG diffuses on dsDNA and that DNA melting is not necessary for the halting of CMG diffusion after ATP-binding. Altogether, our single-molecule and ensemble biochemical data show that the halting of CMG diffusion in the presence of nucleotide is *not* due to DNA melting by CMG, but due to binding of CMG to the DNA *via* other Mcm2-7:DNA interactions^25^.

## Discussion

We report the first single-molecule quantification of the motion of fully reconstituted CMG helicases. To the best of our knowledge, these studies also constitute the first fully *in vitro* reconstituted single-molecule motion quantification of replicative helicases assembled at an origin of replication. To quantify the motion of *de novo* assembled CMG, we developed a novel hybrid ensemble and single-molecule assay based on the double functionalization of DNA ends, which we validated by verifying that the number of fluorescent CMG helicases per diffraction-limited spot, the initial CMG positions on the DNA, and the DDK-dependent colocalization of fluorescently labeled Cdc45 and Mcm2-7 agree with the consensus established by previous biochemical observations (Fig. 1).

In our motion analysis, we observe a static and a mobile population of CMG both in the presence and in the absence of ATP (Fig. 2 and Extended Data Fig. 5). Nonetheless, we quantitatively show that the mobile population in the presence of ATP preferentially moves unidirectionally, as expected for a molecular motor, whereas the mobile population in the absence of ATP preferentially moves in a diffusive manner (Fig. 2). To explain these motion outcomes, we propose the model summarized in Fig. 4c, which is based on the assumption that our ensemble activation reaction contains a mixture of two populations of CMG: one population encircling dsDNA, and another population encircling ssDNA (Fig. 4c-*i*). In the first part of our model, we propose that the population of CMG surrounding ssDNA moves unidirectionally in the presence of ATP (as the motor can then hydrolyze ATP to unwind dsDNA), but remains static in the absence of ATP (as the motor is unable to unwind DNA without ATP hydrolysis^3,6,31^) (Fig. 4c, right half). This part of the model is supported by the fact that we observe very similar instantaneous and average velocities of unidirectionally translocating CMG spots to those of previous single-molecule and ensemble biochemical studies of DNA unwinding^4,5,18^. We consider an alternative scenario in which at least part of the unidirectional translocation that we observe in the presence of ATP corresponds to CMG translocating on dsDNA (Extended Data Fig. 10) – as was postulated to occur for CMG helicases that bypassed each other after the collision of two replication forks or after encountering a flush ss/dsDNA junction^32–34^– to be less likely. Consider, for example, the CMG splitting events that we observe (Fig. 3c). Because CMG translocation on dsDNA has the same 3’-to-5’ polarity as on ssDNA^34^, one could postulate that the splitting events represent two helicases that surround dsDNA following assembly at a single Mcm2-7 double hexamer, but such helicases would move *towards* each other (remaining within a single diffraction-limited spot), and not *away* from each other as our data shows. One could also postulate that the splitting events represent two helicases encircling dsDNA following assembly at distinct Mcm2-7 double hexamers within a single diffraction-limited spot. However, our analysis of Mcm2-7 spots shows such occurrences to be unlikely (Extended Data Fig. 2c). Further studies will be required to assess whether and how the unidirectional motion of CMG differs according to whether it occurs on ssDNA or dsDNA.

In the second part of our model, we propose that the population of CMG surrounding dsDNA loses its bound ATP when we remove ATP from the buffer following our ensemble activation (Fig. 1a); ATP dissociation then causes CMG to disengage from the DNA and thereby become diffusive. This diffusive population remains so in the absence of nucleotide (giving rise to the diffusive population we observe in the absence of nucleotide); nevertheless, upon ATP re-addition and re-binding, CMG is allowed to re-engage with the DNA inside its central channel and thus become static (giving rise to the static population in the presence of ATP). In support of this hypothesis, we showed that CMG can diffuse on dsDNA and that nucleotide binding halts this diffusion (Fig. 4a-b). Furthermore, we show that this halting occurs independent of the previously observed DNA melting by CMG upon nucleotide binding^3,25^ (Extended Data Fig. 9), suggesting that DNA engagement and DNA melting by CMG need not occur concomitantly. We propose that the presence of ATP in the nucleus prevents newly assembled CMG from diffusing along the DNA, poising the helicase to catalyse the initial melting required to initiate replication. Further studies investigating which of the additional Mcm2-7:DNA contacts within CMG^25^ are responsible for halting CMG diffusion upon nucleotide binding will shed further light into our observations.

Our measured diffusion coefficient of freely diffusive CMG under the ionic strength conditions of this study (250 mM K-glutamate) (Extended Data Fig. 5l) is similar to our previously measured diffusion coefficient of single Mcm2-7 hexamers in higher ionic strength conditions (500 mM NaCl)^9^. This observation suggests that CMG diffuses more freely on the DNA than single Mcm2-7 hexamers, which in turn suggests that CMG has fewer contacts with the DNA than Mcm2-7 hexamers as recently confirmed by structural studies^25,35^. We also note that single-molecule studies with pre-formed *D. melanogaster* CMG showed no evidence of CMG diffusion in the presence of ATP^4,5^, in agreement with our observations. However, single-molecule studies with pre-formed *S. cerevisiae* CMG reported extensive diffusive behavior in the presence of ATP^6^. Further studies will be required to investigate the reasons for this discrepancy, and to probe potential differences between fully reconstituted and pre-formed CMG, such as whether the phosphorylation state of Mcm2-7 is different, and whether it could influence the interactions that Mcm2-7, the substrate of DDK^1,13^, have with the DNA.

ATP-dependent switching of motion modes, as we observe here for CMG, has been previously observed to act differently for other protein complexes, such as Type III restriction enzymes^36,37^ and the yeast chromatin remodeler SWR1^38^. In the case of Type III restriction enzymes, ATP hydrolysis triggers diffusion on the DNA^36,37^, whereas in the case of SWR1, ATP binding triggers its diffusion along the DNA^38^. Nonetheless, these behaviors are different than in the case of CMG, whose diffusion is halted and not promoted by nucleotide binding.

Given the complexity and the number of components required to fully reconstitute CMG assembly and activation, *in vitro* single-molecule studies quantifying CMG motion have so far relied on preformed CMG helicases assembled through an unknown mechanism upon overexpression of individual constituents^7,8^. Although these studies have provided us with very important insights into how CMG works, it is unknown whether the assembly mechanism of pre-formed CMG is cell-cycle regulated. It is therefore also unknown whether pre-formed CMG has a similar phosphorylation state to the one it has *in vivo*. In addition, pre-formed CMG requires an artificial region of ssDNA to bind DNA and thus does not allow us to access the intricacies of CMG assembly and activation. In this study, we report the first single-molecule quantification of the motion of fully reconstituted CMG helicases. We believe that the full reconstitution of CMG makes it more likely that the phosphorylation state of its constituents more closely mimics what happens *in vivo*; this in turn makes it more likely that fully reconstituted CMG can support all relevant interactions with proteins intrinsic and accessory to the replisome. Furthermore, the reliable assay we developed will allow us and others to address important questions regarding CMG motion, such as the mechanistic role of elongation factors, with unprecedented resolution. Finally, we anticipate that the novel assay we developed, based on the double functionalization of DNA ends with two orthogonal attachment types, will facilitate the study at the single-molecule level of similarly complex biochemical reactions involved in other nucleic acid:protein interactions.

## Acknowledgements

We thank Anne Early, Lucy Drury, and Max Douglas for providing the yeast strains for the overexpression of unlabeled proteins, Jacob Lewis for providing us with purified M.HpaII methyltransferase, and N.D. lab members Anuj Kumar, Katinka Ligthart, and Julien Gros for assistance in purifying loading proteins and/or DDK. We also thank Kaley McCluskey, Dorian Mikolajczak, Joe Yeeles, Jacob Lewis, and Alessandro Costa for scientific discussions. D.R.M. acknowledges funding from a Boehringer Ingelheim Fonds PhD Fellowship, and N.D. acknowledges funding from the Netherlands Organisation for Scientific Research (NWO) through Top grant 714.017.002, and from the European Research Council through an Advanced Grant (REPLICHROMA; grant number 789267).

## Author contributions

D.R.M., H.S., J.D., and N.D. conceived the study, and N.D. supervised the study. D.R.M. developed the ensemble hybrid and single-molecule study with input from H.S., J.D. and N.D. D.R.M. carried out the ensemble biochemical assays and the did ensemble part of the hybrid ensemble and single-molecule assays. D.R.M. and H.S. acquired the single-molecule data. J.D. provided the strains for the overexpression of unlabeled proteins and advised on biochemical conditions. D.R.M. and T.v.L. carried out the cloning. D.R.M. generated the strains for the overexpression of labeled proteins. D.R.M. and T.v.L. purified the proteins. D.R.M. and H.S. labeled the proteins. E.v.V. designed and wrote data analysis routines with input from all authors as well as conceived and performed the simulations. D.R.M. and E.v.V. performed the data analysis. All authors were involved in the discussion of the data. D.R.M. and N.D. wrote the manuscript with input from all the authors.

## Figure captions

**Extended Data Figure 1.**
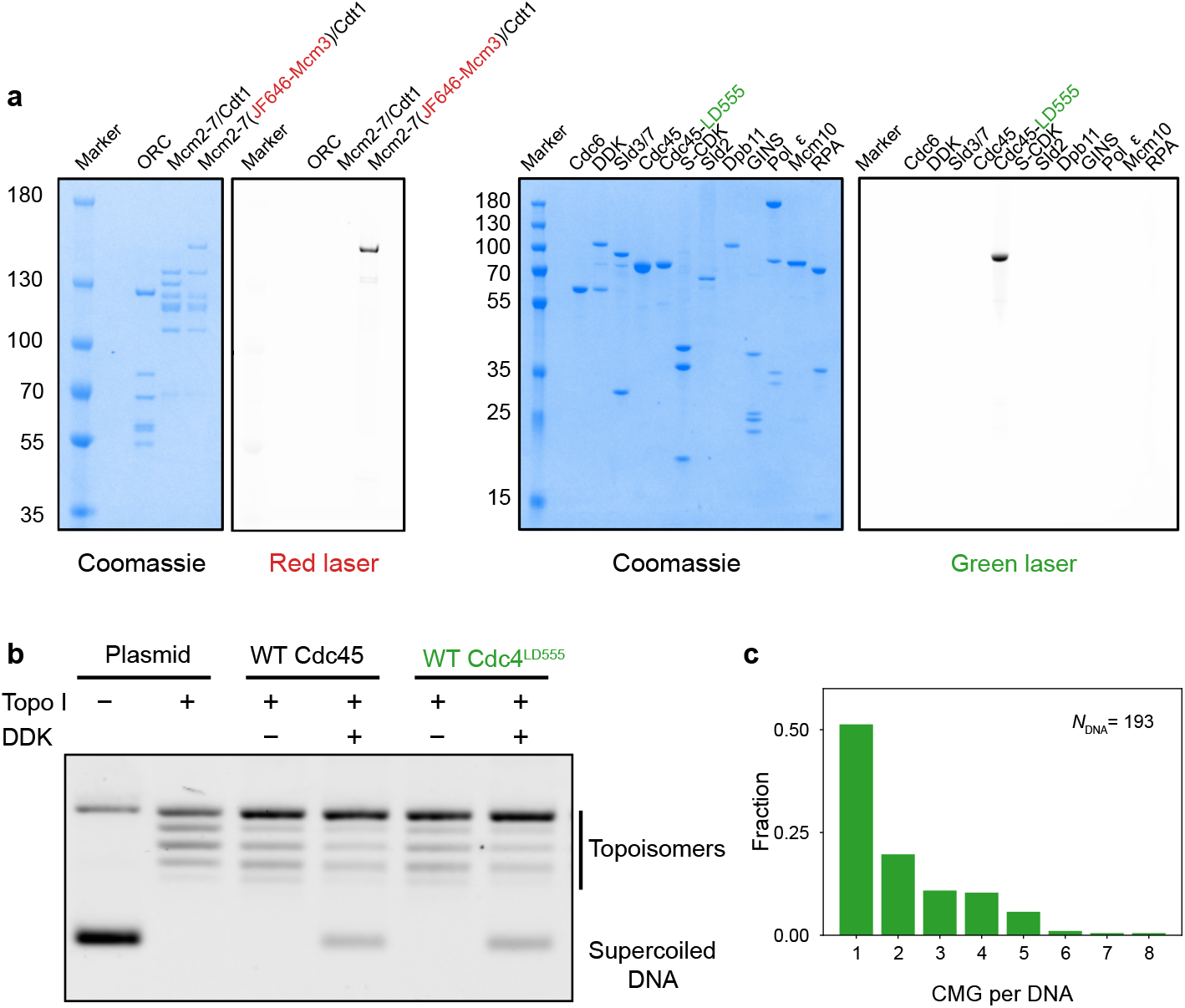
Hybrid ensemble and single-molecule assay and reagent validation. **a**, SDS-PAGE showing the minimal set of purified proteins required for the reconstitution of CMG assembly and activation; the gels were stained with Coomassie Blue Stain and fluorescently scanned with either a red or a green laser, to show the fluorescently labeled proteins in either color. **b**, Ensemble unwinding assay showing that Cdc45^LD555^ supports DNA unwinding to near WT levels. **c**, Distribution of total numbers of fluorescent CMG complexes per DNA, obtained by combining the total number of CMG diffraction-limited spots per DNA (Fig. 1b) with the number of CMG complexes within each spot (Fig. 1c).

**Extended Data Figure 2.**
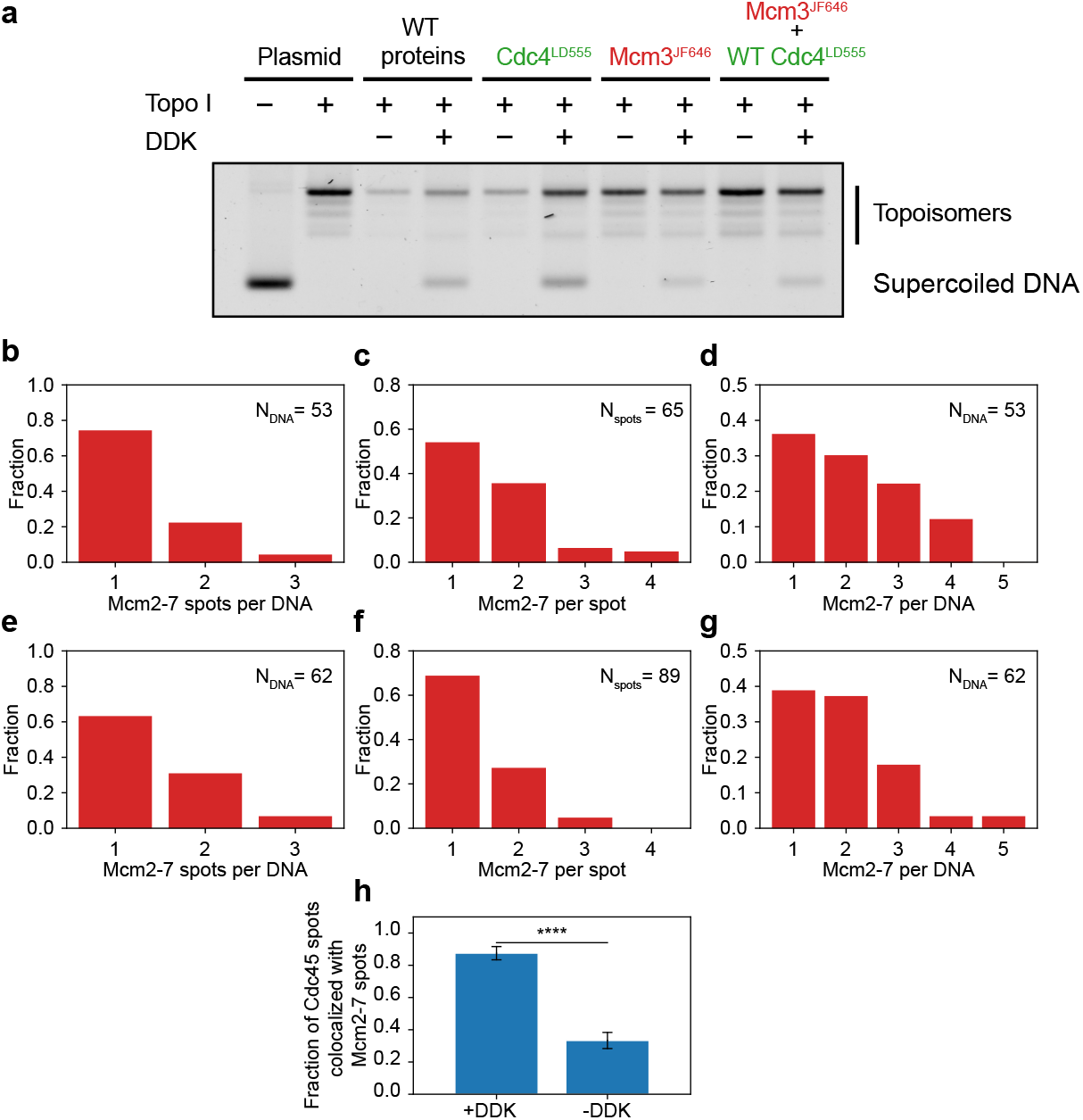
Reagent validation, distribution of number of Mcm2-7 spots and distribution of Mcm2-7 complexes within each spot. **a**, Ensemble unwinding assay showing that Mcm2-7^JF646^ supports DNA unwinding alone and in conjunction with Cdc45^LD555^. **b**, Distribution of the number of Mcm2-7 diffraction-limited spots per DNA in the presence of DDK. **c**, Distribution of the number of Mcm2-7 complexes within each diffraction-limited spot in the presence of DDK. **d**, Distribution of the total number of Mcm2-7 complexes per DNA molecule in the presence of DDK, obtained by combining data from **b** and **c. e**, Distribution of the number of Mcm2-7 diffraction-limited spots per DNA in the absence of DDK. **f**, Distribution of the number of Mcm2-7 complexes within each diffraction-limited spot in the absence of DDK. **g**, Distribution of the total number of Mcm2-7 complexes per DNA molecule in the absence of DDK, obtained by combining data from **e** and **f. h**, Fraction of Cdc45^LD555^ diffraction-limited spots that are colocalized with Mcm2-7^JF646^ diffraction-limited spots in the presence and absence of DDK; error bars show the standard error of proportion. Statistical significance was obtained from a two-sided binomial test (p-value=1.2 × 10^−5^).

**Extended Data Figure 3.**
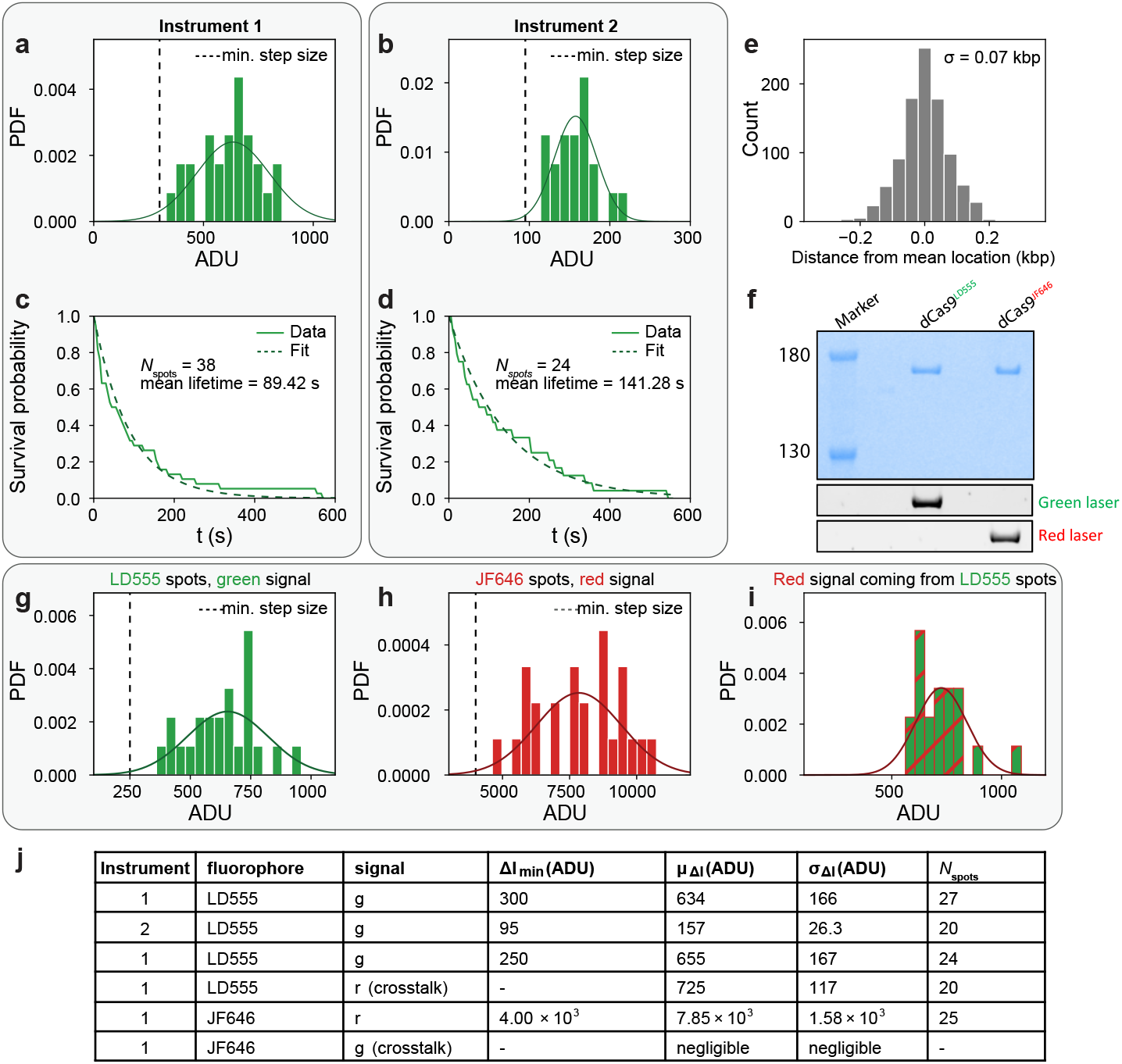
Fluorescently labeled dCas9 proteins as standards for determination of number of proteins per diffraction-limited spot and localization accuracy. **a-b**, Distribution of photobleaching step sizes of fluorescently labeled dCas9^LD555^ imaged under the same imaging conditions as fluorescent CMG in the single-color experiments; **a**, and **b**, correspond to the two instruments used in this study; both distributions were fitted to a normal distribution; μ – 2σ was used as the minimum step size in the single-color CMG experiments to capture at least 95 % of bleaching events. **c-d**, Distribution of times to photobleaching of fluorescently labeled dCas9^LD555^ imaged under the same imaging conditions as fluorescent CMG; **c**, and **d**, correspond to the two instruments used in this study; both distributions were fitted to a single exponential decay. **e**, distribution of positional measurements of fluorescently labeled dCas9^LD555^; as dCas9^LD555^ is expected to be static, the standard deviation of this distribution gives us the localization error in our experiments. **f**, SDS-PAGE of dCas9 with fluorescently labeled with dyes LD555, JF646, respectively; the gel was stained with Coomasie Blue stain and fluorescently scanned with a red, green laser, respectively. **g**, Distribution of photobleaching step sizes of fluorescently labeled dCas9^LD555^ when simultaneously excited with the green and red lasers in instrument 1, as done in the Mcm2-7 and Cdc45 colocalization experiments; the distribution was fitted to a normal distribution; μ – 2σ was used as the minimum step size in the dual-color CMG experiments to capture at least 95 % of bleaching events. **h**, Distribution of photobleaching step sizes of fluorescently labeled dCas9^JF646^ when simultaneously excited with the green and red lasers in instrument 1, as done in the Mcm2-7 and Cdc45 colocalization experiments; the distribution was fitted to a normal distribution; μ – 2σ was used as the minimum step size in the dual-color CMG experiments to capture at least 95 % of bleaching events. **i**, Distribution of red signal coming from green fluorescently labeled dCas9^LD555^ when simultaneously excited with the green and red lasers in instrument 1, as done in the Mcm2-7 and Cdc45 colocalization experiments. The distribution was fitted to a normal distribution and the mean value was used for crosstalk corrections. **j**, Summary table of all the parameters obtained from **a-e**, and **g-i**.

**Extended Data Figure 4.**
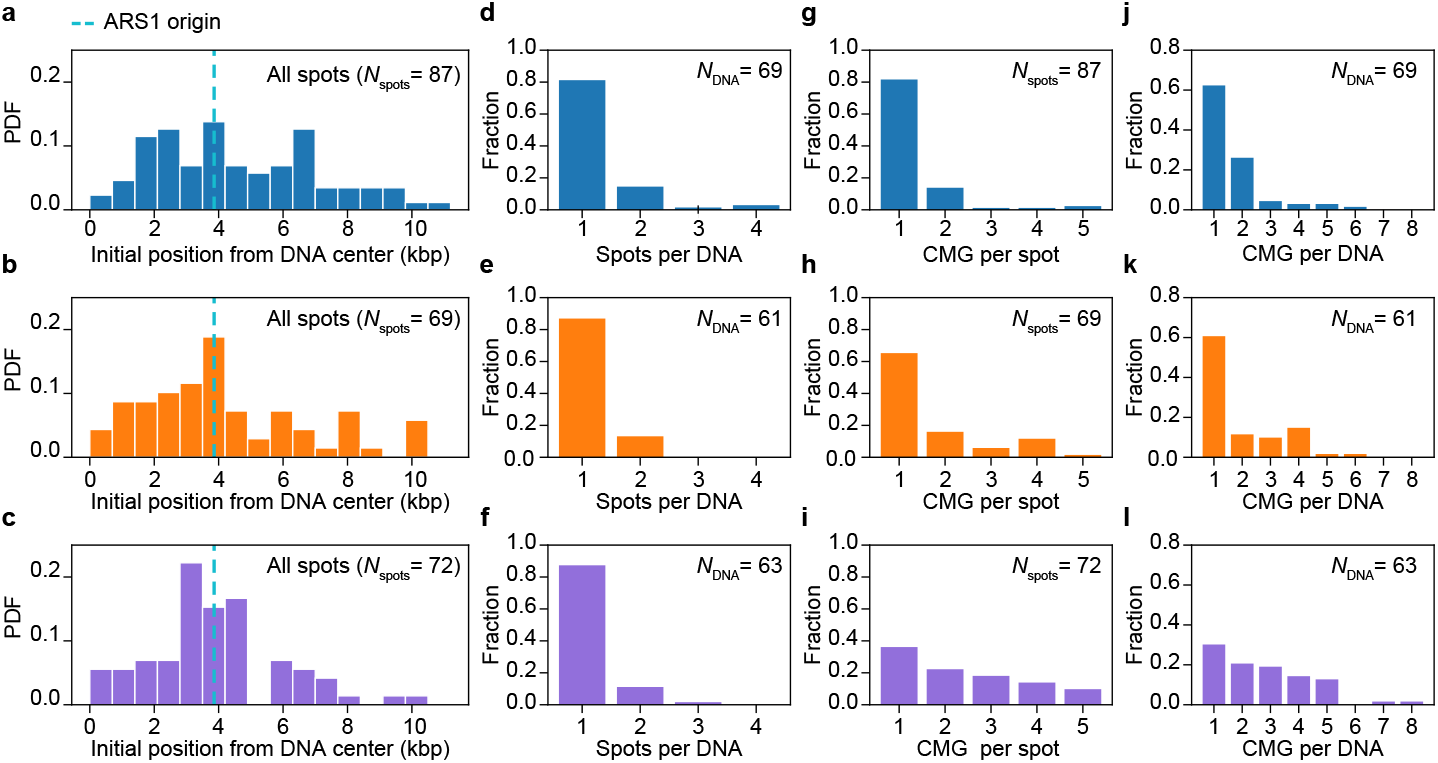
Distribution initial positions, numbers of CMG spots and numbers of CMG complexes within each spot of the different biochemical conditions tested. **a-c**, Distribution of initial positions on the DNA of all Cdc45 diffraction-limited spots for DNA molecules imaged in **a**, the presence of ATP, **b**, the absence of nucleotide or, **c**, the presence of ATPγS. **d-f**, Distribution of numbers of CMG diffraction-limited spots for DNA molecules imaged in **d**, the presence of ATP, **e**, the absence of nucleotide or, **f**, the presence of ATPγS. **g-i**, Distribution of numbers of CMG complexes within each diffraction limited spot on DNA molecules imaged in **g**, the presence of ATP, **h**, the absence of nucleotide or, **i**, the presence of ATPγS. **j-l**, Distribution of numbers of CMG complexes per DNA for DNA molecules imaged in **j**, the presence of ATP, **k**, the absence of nucleotide or, **l**, the presence of ATPγS.

**Extended Data Figure 5.**
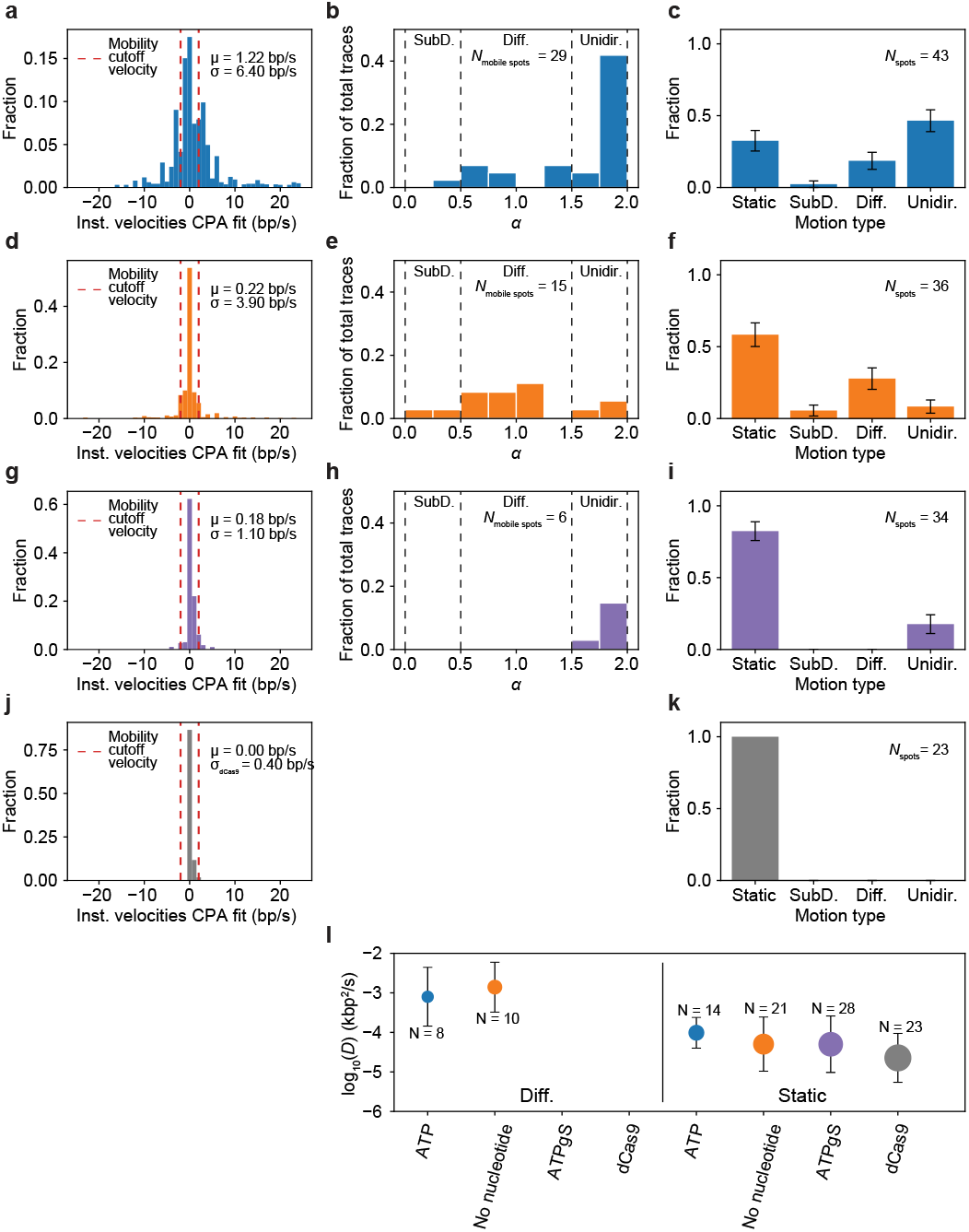
Mobility determination and motion classification of fluorescent spots imaged under different biochemical conditions. **a**, Distribution of instantaneous velocities coming from the CPA fits of CMG spots in the presence of ATP; red lines show the instantaneous velocity cutoff (5σ_dCas9_) used to separate CMG spots into static or mobile. **b**, Distribution of anomalous coefficients α of mobile CMG spots in the presence of ATP. **c**, Fraction of CMG spots imaged in the presence of ATP classified into static, subdiffusive, diffusive or unidirectionally moving. **d**, Distribution of instantaneous velocities coming from the CPA fits of CMG spots in the absence of nucleotide; red lines show the instantaneous velocity cutoff (5σ_dCas9_) used to separate CMG spots into static or mobile. **e**, Distribution of anomalous coefficients α of mobile CMG spots in the absence of nucleotide. **f**, Fraction of CMG spots imaged in the absence of of nucleotide classified into static, subdiffusive, diffusive or unidirectionally moving. **g**, Distribution of instantaneous velocities coming from the CPA fits of CMG spots in the presence of ATPγS; red lines show the instantaneous velocity cutoff (5σ_dCas9_) used to separate CMG spots into static or mobile. **h**, Distribution of anomalous coefficients α of mobile CMG spots in the presence of ATPγS. **i**, Fraction of CMG spots imaged in the presence of ATPγS classified into static, subdiffusive, diffusive or unidirectionally moving. **j**, (*same as inset in Fig. 2a*) Distribution of instantaneous velocities coming from the CPA fits of dCas9^LD555^ spots; red lines show the instantaneous velocity cutoff (5σ_dCas9_) used to separate CMG spots into static or mobile. **k**, Fraction of dCas9^LD555^ spots classified into static, subdiffusive, diffusive or unidirectionally moving. **l**, (left half) Diffusion constants of spots classified as diffusive for the different biochemical conditions tested; (right half) Diffusion constants of spots classified as static for the different biochemical conditions tested.

**Extended Data Figure 6.**
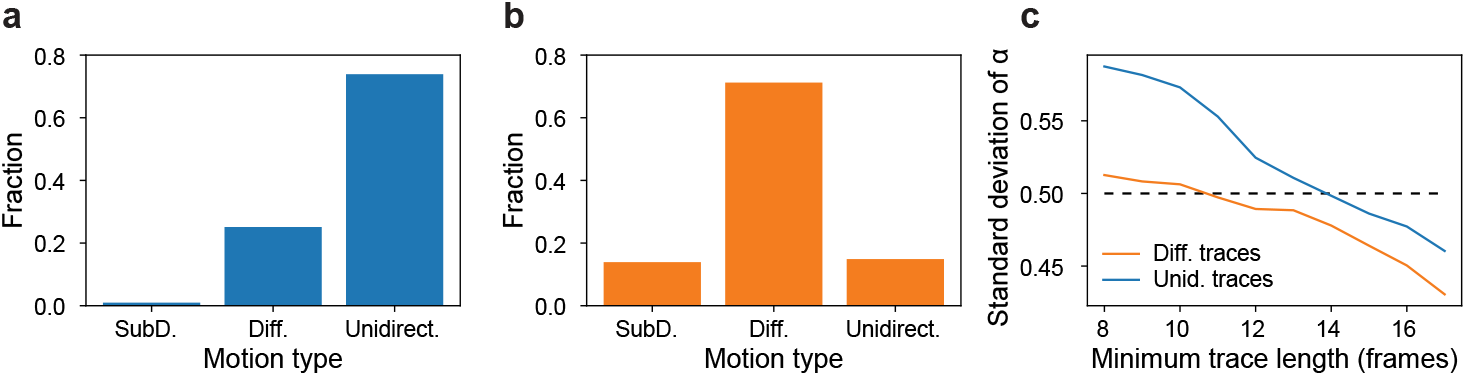
Motion classification of simulated unidirectional or diffusive traces and anomalous diffusion exponent error determination. Motion classification of simulated **a**, unidirectionally translocating traces with a representative velocity (5 bp/s) and **b**, diffusive traces with a representative diffusion coefficient (1.5 × 10^−3^ kb^2^/s). **c**, Error determination of the anomalous diffusion exponent *α* as a function of the minimum trace length; the error falls below 0.5 for a minimum trace length of 14 frames. We start with 512 traces of each motion type with a minimum trace length of 8 pulled from a population with a mean fluorophore lifetime of 25 frames, and gradually increase the trace length filtering. The traces used in **a-b**, are those with a minimum trace length of 14, to mirror the motion analysis done on experimentally obtained CMG spots.

**Extended Data Figure 7.**
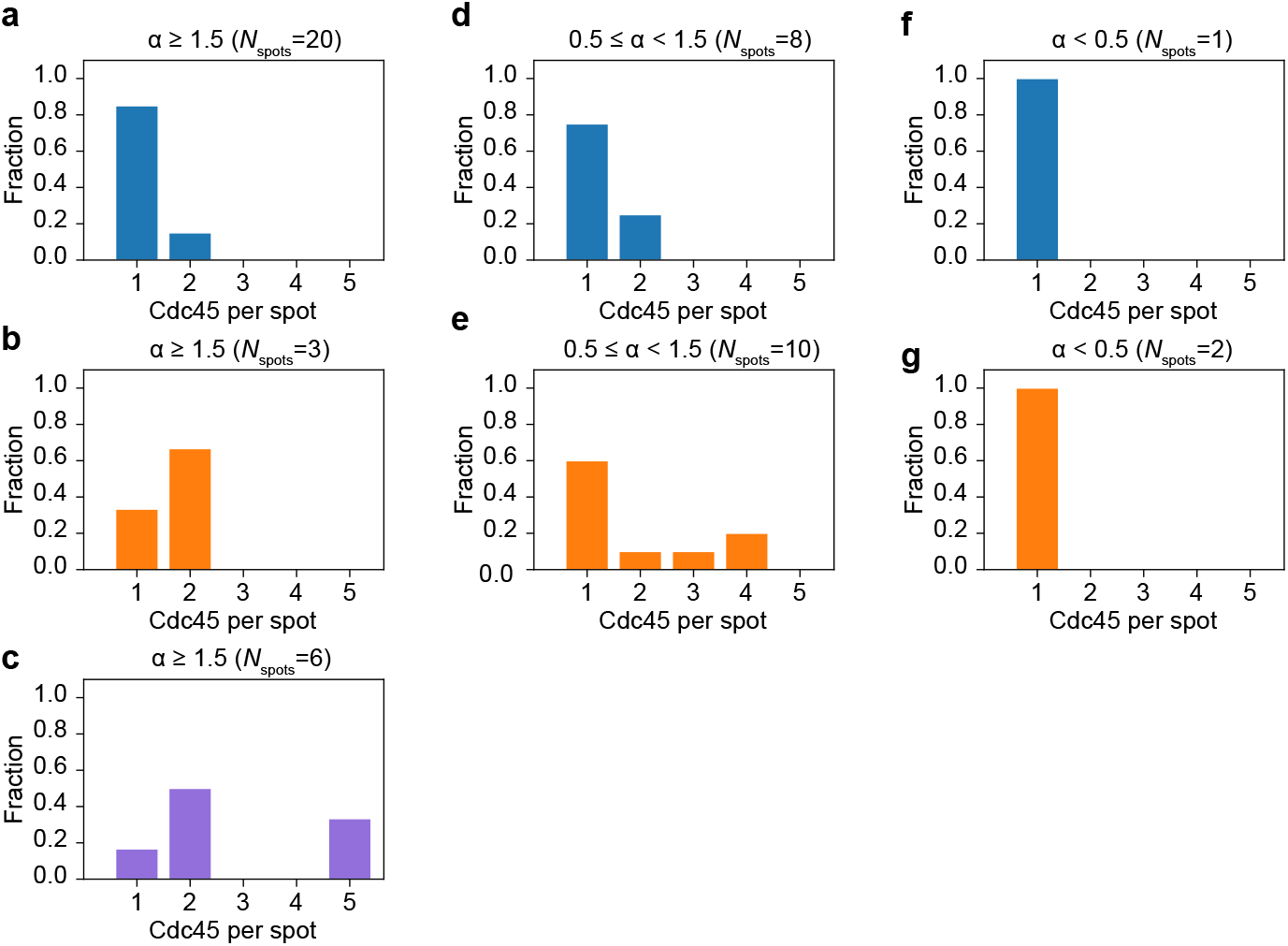
Distribution of number of Cdc45 molecules per mobile diffraction-limited spot for the different biochemical conditions tested. **a-c**, Distribution of number of Cdc45 molecules within diffraction-limited spots classified as unidirectionally-moving in the **a**, presence of ATP, **b**, absence of nucleotide or **c**, presence of ATPγS. **d-e**, Distribution of number of Cdc45 molecules within diffraction-limited spots classified as diffusive in the **d**, presence of ATP or **e**, absence of nucleotide. **f-g**, Distribution of number of Cdc45 molecules within diffraction-limited spots classified as diffusive in the **f**, presence of ATP or **g**, absence of nucleotide.

**Extended Data Figure 8.**
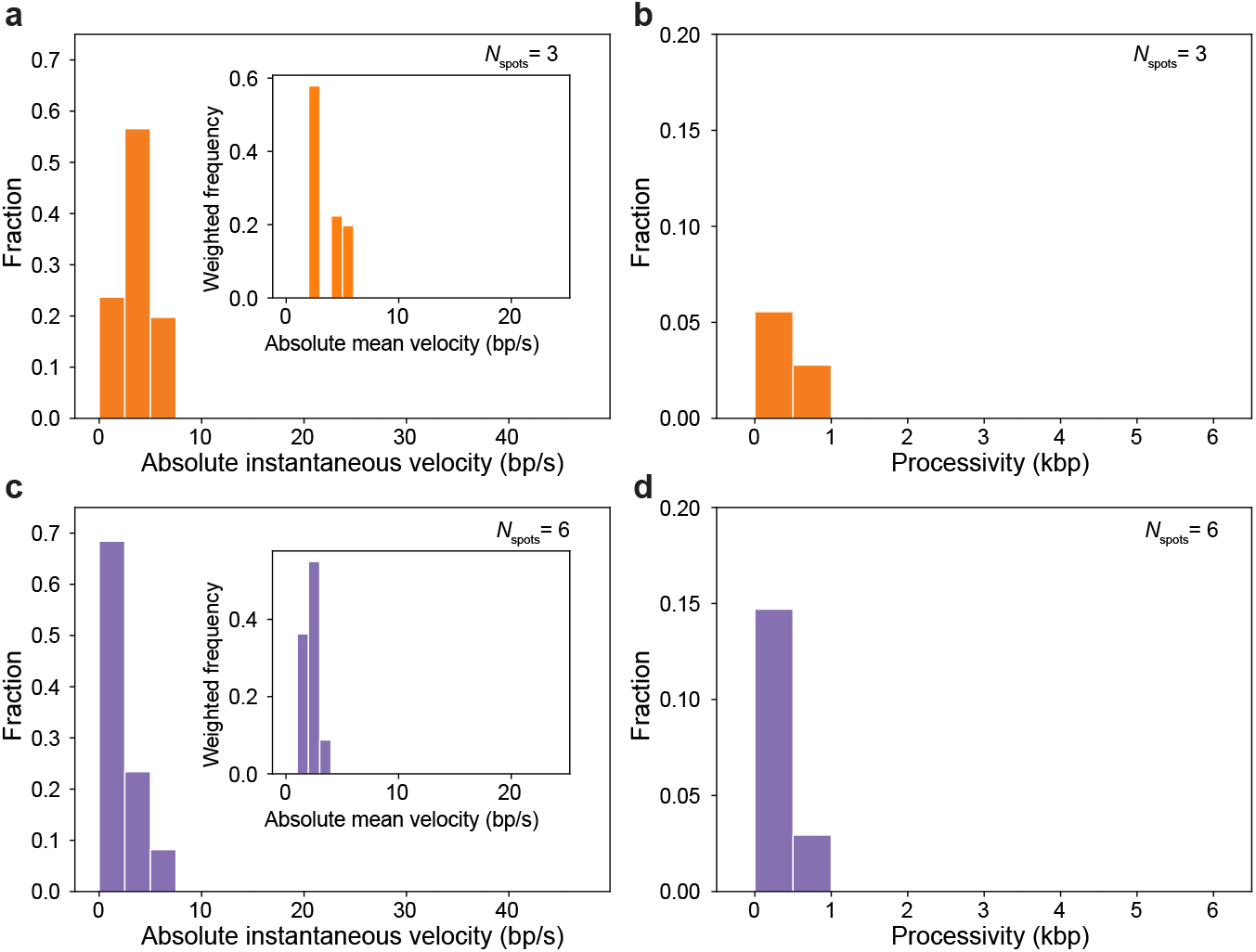
Analysis of unidirectionally moving CMG under different biochemical conditions. **a**, Distribution of absolute instantaneous velocities of unidirectionally moving CMG spots in the absence of nucleotide; (inset) Distribution of absolute mean velocities of unidirectionally moving CMG spots in the absence of nucleotide normalized by the length of each trace. **b**, Distribution of processivities of unidirectionally moving CMG spots in the absence of nucleotide. **c**, Distribution of absolute instantaneous velocities of unidirectionally moving CMG spots in the presence of ATPγS; (inset) Distribution of absolute mean velocities of unidirectionally moving CMG spots in the presence of ATPγS normalized by the length of each trace. **d**, Distribution of processivities of unidirectionally moving CMG spots in the presence of ATPγS.

**Extended Data Figure 9.**
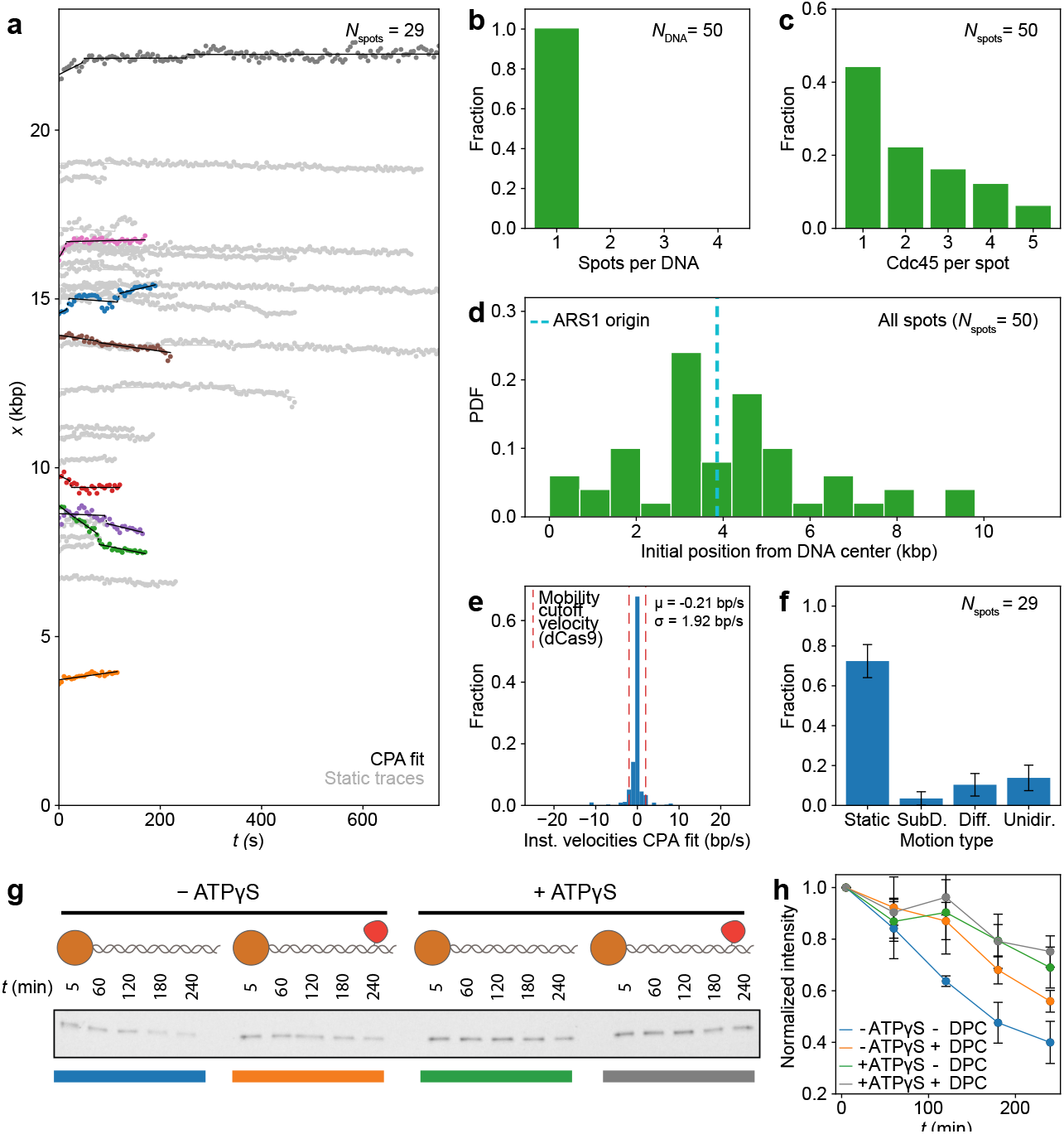
Nucleotide binding halts CMG diffusion independently of DNA melting. **a**, Position *vs*. time plots of CMG^Mcm2(6A)^ spots in the presence of ATP; CPA fits are plotted in black, static traces are shown in light gray and mobile traces are shown in all other colors. **b**, Distribution of numbers of CMG^Mcm2(6A)^ diffraction-limited spots per DNA. **c**, Distribution of numbers of CMG^Mcm2(6A)^ complexes within each diffraction-limited spot. **d**, Distribution of initial positions on the DNA of all CMG^Mcm2(6A)^ diffraction-limited spots. **e**, Distribution of instantaneous velocities coming from the CPA fits of CMG^Mcm2(6A)^ spots in the presence of ATP; red lines show the instantaneous velocity cutoff (5σ_dCas9_) used to separate CMG^Mcm2(6A)^ spots into static or mobile. **f**, Fraction of CMG^Mcm2(6A)^ spots imaged in the presence of ATP classified into static, subdiffusive, diffusive or unidirectionally moving. **g**, Fluorescent scan of an SDS-PAGE gel showing the amount of Cdc45^LD555^ left on linear DNA bound to magnetic beads at one end and containing either a free end or an end capped with a covalently crosslinked methyltransferase. **h**, Densitometry quantification of the experiment shown in **g**, showing the average normalized intensity of three replicates together with their standard deviation. Data points are connected by solid lines to guide the eye.

**Extended Data Figure 10.**
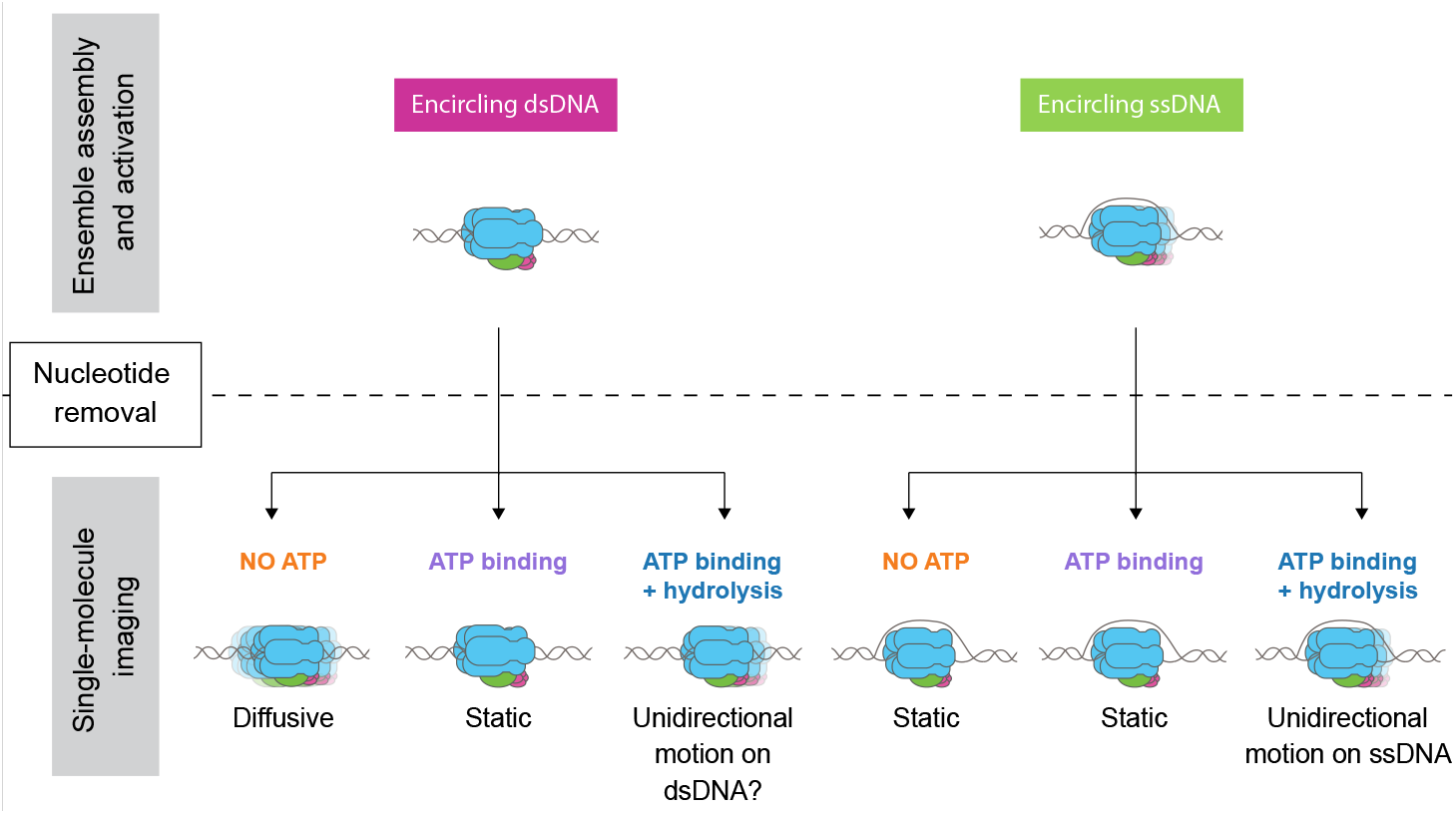
Final model. Model showing all the experimental outcomes observed in this study with different potential explanations.

