## Supplementary Information for "Nucleotide binding halts diffusion of the eukaryotic replicative helicase during activation"

### Methods

#### Biological Materials and Preparation

##### Molecular cloning and strain generation

***Cdc45-iS6-i2XFLAG:*** to generate fluorescently labelled Cdc45, we introduced an 'S6' peptide[39] with an additional glycine linker (GDSLSWLLRLLNG) after residue E197 and immediately before the internal 2XFLAG tag of Cdc45. This was achieved by modifying plasmid pJY13 with a Q5® Site-Directed Mutagenesis Kit (NEB # E0554S) and mutagenic primers DRM_005 and DRM_006. The resulting plasmid pDRM19-01 was transformed into chemically competent One Shot™ TOP10 *E. coli* (Invitrogen # C404010). The sequence of Cdc45-iS6-i2XFLAG was verified by sequencing the entire ORF. *S. cerevisiae* strain yDRM2 overexpressing Cdc45-iS6-i2XFLAG was then generated by linearizing plasmid pDRM19-01 with *NheI*-HF® (NEB # R3131S) and transforming it into yeast strain yJF1 (*MAT***a**, W303 background).

***6XHis-S6-dCas9-Halo:*** to generate fluorescently labelled dCas9, we introduced an 'S6' peptide with an additional GSS linker (GDSLSWLLRLLNGGS) next to the N-terminal 6XHis tag. This was achieved by modifying plasmid pET302-6His-dCas9-halo with Q5® Site-Directed Mutagenesis Kit (NEB # E0554S) and mutagenic primers DRM_184 and DRM_185. The resulting plasmid, pDRM21-01 was transformed into NEB® 5-alpha Competent *E. coli* (High Efficiency) cells (NEB # C2987I). The sequence of pDRM21-01 was then verified by sequencing the entire ORF. Plasmid pDRM21_01 was then transformed into *E. coli* BL21-Codon Plus (DE3)-RIL Competent Cells (Agilent # 230245).

***pGL50-ARS1:*** the ARS1 sequence was amplified by PCR using plasmid p5.8kb-ARS1[2] as a template together with primers TL-033 and TL-034, which contained *Asc*I restriction sites at both ends. This PCR product was then cloned into the dephosphorylated *Mlu*I site of the 21.2 kb plasmid pSupercos1-lambda 1,2[40]. The whole sequence of the ARS1 insert was confirmed by sequencing.

##### Protein purification

Cdc6, Mcm10, dCas9-Halo, S6-dCas9-Halo and Sfp phosphopantetheinyl transferase were expressed in *E. coli* BL21-Codon Plus (DE3)-RIL Competent Cells (Agilent # 230245). GINS was expressed in *E. coli* Bl21 (DE3) Rosetta pLysS Competent Cells (Novagen). Unless otherwise specified, cells were grown to OD600 = 0.40-0.60, induced with 400 $\mu$M IPTG for 16 h at 17 °C, and harvested by centrifugation. Pellets were resuspended in 40 mL of lysis buffer and sonicated on ice in a Qsonica Q500 sonicator for 2 min in 5 s on / 5 s off cycles and an amplitude of 40 %. Following sonification, the lysate was centrifuged at 8820 g for 20 min in a Beckman-Coulter Avanti JXN 26 centrifuge with rotor JA17, and the clarified supernatant was used for affinity pulldowns.

ORC, Mcm2-7/Cdt1, Pol ε, Dpb11, Sld2, Sld3/7, Cdc45, Cdc45-iS6, RPA, DDK, and S-CDK were expressed in *S. cerevisiae* strain yJF1 *MAT***a** Δpep1 Δbar1 from a bi-directional galactose-inducible promotor. Prior to expression, cells were inoculated at a density of 2 x $10^5$ cells/ml in YP medium supplemented with 2 % raffinose (Carbosynth # OR06197) and 50 mg/ml ampicillin (Merck-Sigma # A9518), and grown overnight at 30 °C to a density of 3-5 x $10^7$ cells/ml. For S-CDK, 5 $\mu$g/ml Nocodazole (Merck-Sigma # M1404), and 2 % galactose (Carbosynth # MG05201) were added to the medium and the cells were induced for 3 h before harvesting. For DDK, 2 % galactose (Carbosynth # MG05201) was added to the medium and the cells were induced for 3 h before harvesting. For all other proteins, cells were arrested in G1 phase for 3 h before induction with 100 ng/ml of α mating factor (Tebu-bio # 089AS-60221); cells were then induced with 2 % galactose (Carbosynth # MG05201) for 3 h before harvesting.

Cells were harvested by centrifugation and washed with lysis buffer. After centrifugation, cells were suspended in lysis buffer supplemented with protease inhibitors (cOmpleteTM EDTA-free Protease Inhibitors (Merck-Sigma # 5056489001) and 0.3 mM phenylmethylsulfonyl fluoride (PMSF)), and dropwise dropped into liquid nitrogen. The frozen droplets were grounded in a 6875 SPEX freezer mill for six cycles (run time 2 min and cool time 1 min, with a rate of 15 cps). Following purification, all protein concentrations were determined with Bio-Rad Protein Assay Dye Reagent (Bio-Rad #5000006).

Purifications of ORC, Cdc6, Mcm2-7/Cdt1, Mcm2-7$^{Halo\text{-}Mcm3}$/Cdt1, Mcm2-7$^{Mcm2(6A)}$/Cdt1, and dCas9-Halo have been described previously[9,25].

**DDK:** DDK with a CBP-TEV tag on Dbf4 was purified from *S. cerevisiae* strain ySDK8[41]. Powder was suspended in DDK lysis buffer (25 mM HEPES-KOH pH 7.6. 0.05 % NP40 substitute, 10 % glycerol, 400 mM NaCl, and 1 mM DTT) supplemented with protease inhibitors. Lysate was cleared in a Beckman-Coulter ultracentrifuge (type Optima L90K with rotorTI45) for 1 h at 45000 rpm and 4 °C. The cleared lysate was supplemented with $CaCl_2$ to a final concentration of 2 mM and incubated for 1 h at 4 °C with washed Sepharose 4B Calmodulin beads (GE Healthcare # 17-0529-01) in a spinning rotor. The beads were washed 10 times with 5 ml of DDK-binding buffer (25 mM HEPES-KOH pH 7.6, 0.05% NP40 substitute, 10% glycerol, 400 mM NaCl, 2 mM $CaCl_2$, and 1 mM DTT), and protein was eluted from the beads with DDK elution buffer (25 mM HEPES-KOH pH 7.6, 0.05 % NP40 substitute, 10 % glycerol, 400 mM NaCl, 2 mM EDTA, 2mM EGTA, and 1 mM DTT). DDK-containing fractions were pooled and dephosphorylated with 20000 units lambda phosphatase (NEB # P0753S) for 16 h at 4°C. The sample was then concentrated in an Amicon Ultra-4 Ultracell 30 kDa centrifugal filter (Merck-Millipore # UFC803024), and injected into a Superdex 200 Increase 10/300 GL column (Cytiva # 15182085) equilibrated in DDK GF buffer (25 mM HEPES-KOH pH 7.6, 0.02 % NP40 substitute, 10 % glycerol, and 200 mM K glutamate). Positive fractions were pooled and concentrated in an Amicon Ultra-4 Ultracell 30 kDa centrifugal filter (Merck-Millipore # UFC803024). Aliquots were snap frozen and stored at −80 °C.

**GINS:** GINS with a N-terminal 6xHis-tag on Psf3 was purified from *E. coli Rosetta*. Cleared lysate was incubated with Ni-NTA agarose (Qiagen # 30210) for 1 h at 4 °C in GINS lysis buffer (25 mM Tris-HCl pH 7.2, 0.02 % NP40 substitute, 10 % glycerol, 400 mM NaCl, 10 mM imidazole, and 1 mM DTT) supplemented with protease inhibitors. The beads were washed 5 times with 5 ml of GINS lysis buffer and 5 times with 5 ml of GINS wash buffer (25 mM Tris-HCl pH 7.2, 0.02 % NP40 substitute, 10 % glycerol, 100 mM NaCl, 15 mM imidazole and 1 mM DTT). Protein was then eluted from the beads with GINS elution buffer (25 mM Tris-HCl pH 7.2, 0.02 % NP40 substitute, 10 % glycerol, 100 mM NaCl, 200 mM imidazole, and 1 mM DTT). GINS-containing fractions were pooled and dialyzed against GINS dialysis buffer I (25 mM Tris-HCl pH 7.2, 0.02 % NP40 substitute, 10 % glycerol, 100 mM NaCl, and 1 mM DTT). The sample was then flowed through a 0.2 micron filter and injected into a MonoQ 5/50 GL column (GE Healthcare # 17-5166-01) equilibrated in GINS MonoQ buffer A (25 mM Tris-HCl pH 7.2, 0.02 % NP40 substitute, 10 % glycerol, 100 mM NaCl, and 1 mM DTT). GINS was eluted from the column in a 30 CV NaCl gradient from 100 mM to 500 mM. Positive fractions were pooled, concentrated in an Amicon Ultra-4 Ultracell 30 kDa centrifugal filter (Merck-Millipore # UFC803024), and injected into a Superdex 200 Increase 10/300 GL column (Cytiva # 15182085) equilibrated in GINS GF buffer (25 mM Tris-HCl pH 7.2, 0.02 % NP40 substitute, 10 % glycerol, 150 mM NaCl, and 1 mM DTT). Positive fractions were pooled and concentrated in an Amicon Ultra-4 Ultracell 30 kDa centrifugal filter (Merck-Millipore # UFC803024). Aliquots were snap frozen and stored at −80°C.

**Pol ε:** Pol ε with a C-terminal CBP-TEV tag on Dpb4 was purified from *S. cerevisiae* strain yAJ2[2]. Powder was suspended in Pol ε lysis buffer (25 mM HEPES-KOH pH 7.6, 10 % glycerol, 400 mM KOAc, and 1 mM DTT) supplemented with protease inhibitors. Lysate was cleared in a Beckman-Coulter ultracentrifuge (type Optima L90K with rotor TI45) for 1 h at 45000 rpm and 4 °C. The cleared lysate was supplemented with $CaCl_2$ to a final concentration of 2 mM and incubated for 1 h at 4°C with washed Sepharose 4B Calmodulin beads (GE Healthcare # 17-0529-01) in a spinning rotor. The beads were washed 10 times with 5 ml of Pol ε binding buffer (25 mM HEPES-KOH pH 7.6, 10 % glycerol, 400 mM KOAc, 2 mM $CaCl_2$, and 1 mM DTT) and protein was eluted from the beads with Pol ε elution buffer (25 mM HEPES-KOH pH 7.6, 10 % glycerol, 400 mM KOAc, 2 mM EDTA, 2 mM EGTA, and 1 mM DTT). Positive fractions were pooled, passed through a 0.2 micron filter and injected into a 1-ml heparin column (GE Healthcare # 17-0406-01) equilibrated in Pol ε heparin buffer A1 (25 mM HEPES-KOH pH 7.6, 10 % glycerol, 400 mM KOAc, and 1 mM DTT). The column was washed with Pol ε heparin buffer A2 (25 mM HEPES-KOH pH 7.6, 10 % glycerol, 450 mM KOAc, and 1 mM DTT) and Pol ε was eluted from the column in a 30 CV KOAc gradient from 450 mM to 1000 mM. Peak fractions were pooled, concentrated in an Amicon Ultra-4 Ultracell 30 kDa centrifugal filter (Merck-Millipore # UFC803024), and injected into a Superdex 200 Increase 10/300 GL column (Cytiva # 15182085) equilibrated in Pol ε GF buffer (25 mM HEPES-KOH pH 7.6, 10 % glycerol, 500 mM KOAc, and 1 mM DTT). Peak fractions were pooled and concentrated in an Amicon Ultra-4 Ultracell 30 kDa centrifugal filter (Merck-Millipore # UFC803024). Aliquots were snap frozen and stored at –80°C.

**S-CDK**: S-CDK with an N-terminal CBP-TEV tag on Clb5 Δ1-100 was purified from *S. cerevisiae* (Lucy Drury, Francis Crick Institute, unpublished). Powder was suspended in S-CDK lysis buffer (40 mM HEPES-KOH pH 7.6, 0.01 % NP40 substitute, 10 % glycerol, and 300 mM KOAc) supplemented with protease inhibitors. The lysate was cleared in a Beckman-Coulter ultracentrifuge (type Optima L90K with rotorTI45) for 1 h at 45000 rpm and 4°C. The cleared lysate was supplemented with $CaCl_2$ to a final concentration of 2 mM and incubated for 1 h at 4 °C with washed Sepharose 4B Calmodulin beads (Agilent # 214303-52) in a spinning rotor. The beads were washed 5 times with 5 ml of S-CDK binding buffer (40 mM HEPES-KOH pH 7.6, 0.02 % NP40 substitute, 10 % glycerol, 300 mM KOAc, and 2 mM $CaCl_2$), and 5 times with 5 ml of S-CDK-TEV buffer (40 mM HEPES-KOH pH 7.6, 0.02 % NP40 substitute, 10 % glycerol, and 2 mM $CaCl_2$). S-CDK was then cleaved from the beads by incubating overnight with TEV protease at 4 °C in S-CDK TEV buffer. Released S-CDK was then concentrated in an Amicon Ultra-4 Ultracell 30 kDa centrifugal filter (Merck-Millipore # UFC803024), and injected into a Superdex 200 Increase 10/300 GL column (Cytiva # 15182085) equilibrated in S-CDK GF buffer (40 mM HEPES-KOH pH 7.6, 0.01 % NP40 substitute, 10 % glycerol, and 300 mM KOAc). Peak fractions were pooled and concentrated in an Amicon Ultra-4 Ultracell 30 kDa centrifugal filter (Merck-Millipore # UFC803024). Aliquots were snap frozen and stored at –80 °C.

**Dpb11**: Dpb11 with a C-terminal 3xFlag-tag was purified from S. *cerevisiae* strain yJY26[2]. Powder was suspended in Dpb11 lysis buffer (25 mM HEPES-KOH pH 7.6, 0.02 % NP40 substitute, 10 % glycerol, 500 mM KCl, 1 mM EDTA, and 1 mM DTT) supplemented with protease inhibitors. The lysate was cleared in a Beckman-Coulter ultracentrifuge (type Optima L90K with rotor TI45) for 1 h at 45000 rpm and 4 °C. The cleared lysate was incubated for 1 h at 4 °C with washed M2 anti-flag affinity beads (Merck-Sigma # A2220) in a spinning rotor. The beads were washed 10 times with 5 ml Dpb11 lysis buffer and Dpb11 was eluted from the beads by incubation with 3xFLAG peptides (Merck-Sigma # F4799). The eluate was dialysed against Dpb11 dialysis buffer I (25 mM HEPES-KOH pH 7.6, 0.02 % NP40 substitute, 10 % glycerol, 150 mM KCl, 1 mM EDTA, and 1 mM DTT), passed through a 0.2-micron filter and injected into a 1-ml MonoS 5/50 GL column (GE Healthcare # 17-5168-01) equilibrated in Dpb11 MonoS buffer A (25 mM HEPES-KOH pH 7.6. 0.02 % NP40 substitute, 10 % glycerol, 150 mM KCl, 1 mM EDTA, and 1 mM DTT). The column was washed with Dpb11 MonoS buffer A, and Dpb11 was eluted from the column in a 20 CV KCl gradient from 150 mM to 1000 mM. Peak fractions were pooled, concentrated in an Amicon Ultra-4 Ultracell 30 kDa centrifugal filter (Merck-Millipore # UFC803024), and injected into a Superdex 200 Increase 10/300 GL column (Cytiva # 15182085) equilibrated in Dpb11 GF buffer (25 mM HEPES-KOH pH 7.6, 0.02 % NP40 substitute, 10 % glycerol, 300 mM KOAc, 1 mM EDTA, and 1 mM DTT). Peak fractions were pooled and concentrated in an Amicon Ultra-4 Ultracell 30 kDa centrifugal filter (Merck-Millipore # UFC803024). Aliquots were snap frozen and stored at –80 °C.

**Sld2**: Sld2 with a C-terminal 3xFLAG-tag was purified from *S. cerevisiae* strain yTD8[2]. Powder was suspended in Sld2 lysis buffer (25 mM HEPES-KOH pH 7.6, 0.02 % NP40 substitute, 10 % glycerol, 500 mM KCl, 1 mM EDTA, and 1 mM DTT) supplemented with 0.3 mM PMSF, 7.5 mM benzamidine,

0.5 mM AEBSF, 1 mM leupeptin, 1 mM pepstatin A, and 1 μg/ml aprotinin. The lysate was cleared in a Beckman-Coulter ultracentrifuge (type Optima L90K with rotorTI45) for 1 h at 45000 rpm and 4 °C. Solid ammonium sulphate was added to the cleared lysate up to a saturation of 32 %. After 15 min of tumbling at 4 °C, the lysate was cleared by centrifugation at 27000 g for 20 min. Then, solid ammonium sulphate was added to the supernatant up to a saturation of 48 %. After 15 min of tumbling at 4 °C, the lysate was cleared by centrifugation at 27000 g for 20 min. The pellet was dissolved in Sld2 lysis buffer supplemented with 0.3 mM PMSF, 7.5 mM benzamidine, 0.5 mM AEBSF, 1 mM leupeptin, 1 mM pepstatin A, and 1 μg/ml aprotinin, and incubated for 30 min at 4 °C with washed M2 anti-FLAG beads (Merck-Sigma # A2220) in a spinning rotor. The beads were washed 8 times with 5 ml of Sld2 lysis buffer, tumbled for 10 min at 4 °C with 10 ml of FLAG resuspension buffer (25 mM HEPES-KOH pH 7.6, 0.02 % NP40 substitute, 10 % glycerol, 500 mM KCl, 1 mM ATP, 10 mM MgOAc, and 1 mM DTT) supplemented with 0.3 mM PMSF, 7.5 mM benzamidine, 0.5 mM AEBSF, 1 mM leupeptin, 1 mM pepstatin A, and 1 μg/ml aprotinin, and 8 more times with 5 ml of Sld2 lysis buffer. Sld2 was eluted from the beads by incubation with 3xFLAG peptides (Merck-Sigma # F4799). The eluate was dialyzed for 45 min at 4 °C against Sld2 dialysis buffer I (25 mM HEPES-KOH pH 7.6, 0.02 % NP40 substitute, 10 % glycerol, 280 mM KCl, 1 mM EDTA, and 1 mM DTT), passed through a 0.2 micron filter, and injected into a 1-ml HiTrap SPFF column (GE Healthcare # 17-5054-01) equilibrated in Sld2 SPFF buffer A (25 mM HEPES-KOH pH 7.6, 0.02 % NP40 substitute, 10 % glycerol, 250 mM KCl, 1 mM EDTA, and 1 mM DTT). The column was washed with Sld2 SPFF buffer A, and Sld2 was eluted from the column in a 20-CV KCl gradient from 280 mM to 1000 mM. Peak fractions were pooled, dialysed for 45 min at 4 °C against Sld2 dialysis buffer II (25 mM HEPES-KOH pH 7.6, 0.02 % NP40 substitute, 40 % glycerol, 350 mM KCl, 1 mM EDTA, and 1 mM DTT). Aliquots were snap frozen and stored at −80 °C.

**Sld3/7**: Sld3/7 with a C-terminal CTP tag on Sld3 was purified from *S. cerevisiae* strain yTD6[2]. Powder was suspended in Sld3/7 lysis buffer (25 mM HEPES-KOH pH 7.6, 0.02 % NP40 substitute, 10 % glycerol, 500 mM KCl, 1 mM EDTA, and 1 mM DTT) supplemented with protease inhibitors. The lysate was cleared in a Beckman-Coulter ultracentrifuge (type Optima L90K with rotor TI45) for 1 h at 45000 rpm and 4 °C, and incubated for 40 min at 4 °C with washed IgG Sepharose 6 Fast Flow (GE Healthcare cat # 17-0969-01) in a spinning rotor. The beads were washed with 15CV Sld3/7 wash buffer (25 mM HEPES-KOH pH 7.6, 0.02 % NP40 substitute, 10 % glycerol, 500 mM KCl, 0,5 mM EDTA, and 1 mM DTT), and the protein complex was cleaved from the beads by overnight treatment at 4 °C with TEV protease in Sld3/7 lysis buffer. Sld3/7 was eluted from the column, concentrated in an Amicon Ultra-4 Ultracell 30 kDa centrifugal filter (Merck-Millipore # UFC803024), and injected into a Superdex 200 Increase 10/300 GL column (Cytiva # 15182085) equilibrated in Sld3/7 GF buffer (25 mM hepes-KOH pH 7.6, 0.02 % NP40 substitute, 10 % glycerol, 500 mM KCl, 1 mM EDTA, and 1 mM DTT). Peak fractions were pooled and concentrated in an Amicon Ultra-4 Ultracell 30 kDa centrifugal filter (Merck-Millipore # UFC803024). Aliquots were snap frozen and stored at −80 °C.

**Cdc45 and Cdc45-iS6**: Cdc45 with an internal 2xFLAG tag and Cdc45 with an internal 2xFLAG + S6 tag were purified from S. *cerevisiae* strains yJY13[2] and yDRM2 (this study), respectively. Powder was suspended in Cdc45 lysis buffer (25 mM HEPES-KOH pH 7.6, 10 % glycerol, 500 mM KOAc, 1 mM EDTA, and 1 mM DTT) supplemented with protease inhibitors. The lysate was cleared in a Beckman-Coulter ultracentrifuge (type Optima L90K with rotor TI45) for 1 h at 45000 rpm and 4 °C. The cleared lysate was incubated for 1 h with washed M2 anti-flag affinity beads (Merck-Sigma # A2220) at 4 °C in a spinning rotor. The beads were washed 10 times with 5 ml of Cdc45 lysis buffer, and Cdc45 was eluted from the beads by incubation with 3xFLAG peptides (Merck-Sigma # F4799). The eluate was dialyzed against Cdc45 dialysis buffer I (20 mM K phosphate pH 7.6, 10 % glycerol, 150 mM KOAc, and 0.5 mM DTT) and injected into a 2-ml hydroxyapatite Bio gel HTP column (Biorad # 130-0420) equilibrated in Cdc45 HTP equilibration buffer (20 mM K phosphate pH 7.6, 10 % glycerol, 150 mM KOAc, and 0.5 mM DTT). Cdc45 bound to the hydroxyapatite Bio gel for 45 min at 4 °C with tumbling. The column was then washed with Cdc45 wash buffer A (80 mM K phosphate pH 7.6, 10 % glycerol, 150 mM KOAc, and 0.5 mM DTT), and Cdc45 was eluted with Cdc45 HTP elution buffer (250 mM K phosphate pH 7.6, 10 % glycerol, 150 mM KOAc and 0.5 mM DTT). Positive fractions were pooled and dialyzed against Cdc45 dialysis buffer I (25 mM HEPES-KOH pH 7.6, 10 % glycerol, 300 mM KOAc, 1 mM EDTA, and 1 mM DTT). Finally, Cdc45 was concentrated in an Amicon Ultra-4 Ultracell 30 kDa centrifugal filter (Merck-Millipore # UFC803024). Aliquots were snap frozen and stored at −80 °C.

**RPA:** RPA with a CBP-TEV tag on Rfa1 was purified from *S. cerevisiae* strain yAE31[2]. Powder was suspended in RPA lysis buffer (25 mM Tris-HCl pH 7.2, 10 % glycerol, 500 mM NaCl, and 1 mM DTT) supplemented with protease inhibitors. The lysate was cleared in a Beckman-Coulter ultracentrifuge (type Optima L90K with rotorTI45) for 1 h at 45000 rpm and 4 °C. The cleared lysate was diluted with an equal volume of RPA dilution buffer I (25 mM Tris-HCl pH 7.2, 10% glycerol, and 1 mM DTT), supplemented with $CaCl_2$ to a final concentration of 2 mM, and incubated for 1 h at 4 °C with washed Sepharose 4B Calmodulin beads (GE Healthcare # 17-0529-01) in a spinning rotor. The beads were washed 10 times with 5 ml of RPA binding buffer (25 mM Tris-HCl pH 7.2, 10 % glycerol, 200 mM NaCl, 2 mM $CaCl_2$, and 1 mM DTT), and the protein complex was eluted from the beads with RPA elution buffer (25 mM Tris-HCl pH 7.2, 10 % glycerol, 200 mM NaCl, 2 mM EDTA, 2mM EGTA, and 1 mM DTT). Positive fractions were pooled and diluted with an equal volume of RPA dilution buffer II (25 mM Tris-HCl pH 7.2, 10 % glycerol, 1 mM EDTA, and 1 mM DTT) and twice dialyzed for 1 h against RPA dialysis buffer (25 mM Tris-HCl pH 7.2, 10 % glycerol, 50 mM NaCl, 1 mM EDTA, and 1 mM DTT). The sample was then loaded onto a 1-ml Hi Trap heparin HP column (GE Healthcare # 17-0406-01) equilibrated with buffer RPA heparin A (25 mM Tris-HCl pH 7.2, 10 % glycerol, 50 mM NaCl, 1 mM EDTA, and 1 mM DTT). The column was washed with RPA heparin buffer A, and RPA was eluted from the column in a 30 CV NaCl gradient from 50 mM to 1000 mM. Subsequently, the sample was concentrated in an Amicon Ultra-4 Ultracell 30 kDa centrifugal filter (Merck-Millipore # UFC803024), and injected into a Superdex 200 Increase 10/300 GL column (Cytiva # 15182085) equilibrated in RPA GF buffer (25 mM Tris-HCl pH 7.2, 10 % glycerol, 150 mM NaCl, 1 mM EDTA, and 1 mM DTT). Peak fractions were pooled and concentrated in an Amicon Ultra-4 Ultracell 30 kDa centrifugal filter (Merck-Millipore # UFC803024). Aliquots were snap frozen and stored at −80 °C.

**Mcm10:** Mcm10 with a N-terminal 6xHis-tag and a C-terminal 3xFLAG tag (Max Douglas, Francis Crick Institute, unpublished) was purified from *E. coli* BL21-Codon Plus (DE3)-RIL. Cleared lysate was incubated for 1 h at 4 °C with washed M2 anti-flag affinity beads (Merck-Sigma # A2220) in a spinning rotor. The beads were washed 10 times with 5 ml of Mcm10 lysis buffer (25 mM Tris-HCl pH 7.2, 10 % glycerol, 500 mM NaCl, and 0,01 % NP40 substitute) and 5 times with 5 ml of Mcm10 lysis buffer with 300 mM NaCl (25 mM Tris-HCl pH 7.2, 10 % glycerol, 300 mM NaCl, and 0,01 % NP40 substitute). Mcm10 was eluted from the beads by incubation with 3xFLAG peptides (Merck-Sigma # F4799) and incubated with Ni-NTA agarose (Qiagen # 30210) for 1 h at 4 °C. The beads were washed 5 times with 5 ml of Mcm10 wash buffer II (25 mM Tris-HCl pH 7.2, 10 % glycerol, 500 mM NaCl, and 0,01 % NP40 substitute) and 5 times with 5 ml of Mcm10 wash buffer III (25 mM Tris-HCl pH 7.2, 10 % glycerol, 500 mM NaCl, 0,01 % NP40 substitute, and 20 mM Imidazole). Then the protein complex was eluted from the beads with RPA elution buffer (25 mM Tris-HCl pH 7.2, 10 % glycerol, 500 mM NaCl, 0,01 % NP40 substitute, and 200 mM Imidazole). Positive fractions were pooled and dialyzed against Mcm10 dialysis buffer (25 mM HEPES-KOH pH 7.6, 10 % glycerol, 200 mM KOAc, 1 mM EDTA, and 0.01 % NP40 substitute). After dialysis, Mcm10 was concentrated in an Amicon Ultra-4 Ultracell 30 kDa centrifugal filter (Merck-Millipore # UFC803024). Aliquots were snap frozen and stored at −80 °C.

**S6-dCas9-Halo**: S6-dCas9-Halo with an N-terminal 6xHis-tag (this study) was purified from *E. coli* BL21-Codon Plus (DE3)-RIL. Cleared lysate was incubated with Ni-NTA agarose (Qiagen # 30210) for 1 h at 4 °C in Cas9 lysis buffer (50 mM Na-phosphate pH 7.0, and 300 mM NaCl) supplemented with protease inhibitors. Beads were washed 5 times with 5 ml of dCas9 wash buffer I (50 mM Na phosphate pH 7.0, 300 mM NaCl, and 20 mM imidazole). Then the protein complex was eluted from the beads with dCas9 elution buffer I (50 mM Na phosphate pH 7.0, 300 mM NaCl, and 150 mM imidazole). Positive fractions were pooled and dialyzed against dCas9 dialysis buffer I (50 mM HEPES-KOH pH 7.6, 100 mM KCl, and 1 mM DTT). Then the sample was passed through a 0.2 micron filter and injected into a 1-ml HiTrap SP HP column (GE Healthcare # 17-1151-01) equilibrated in dCas9 SPHP buffer A (50 mM HEPES-KOH pH 7.6, 100 mM KCl, and 1 mM DTT). S6-dCas9 was eluted from the column in a 30 CV KCl gradient from 100 mM to 1000 mM. Positive fractions were concentrated in an Amicon Ultra-4 Ultracell 30 kDa centrifugal filter (Merck Millipore # UFC803024), and injected into a Superose 6 Increase 10/300 GL column (GE Healthcare # 29-0915-96) equilibrated in dCas9 GF buffer (50 mM HEPES-KOH pH 7.6, 150 mM KCl, 20 % glycerol, and 1 mM DTT). Peak fractions were pooled and concentrated in an Amicon Ultra-4 Ultracell 30 kDa centrifugal filter (Merck-Millipore # UFC803024). Aliquots were snap frozen and stored at −80 °C.

**Sfp phosphopantetheinyl transferase:** Sfp phosphopantetheinyl transferase with a C-terminal 6xHis-tag (Addgene # 75015) was purified from *E. coli* BL21-Codon Plus (DE3)-RIL. Cleared lysate was incubated with Ni-NTA agarose (Qiagen # 30210) for 30 min at 4 ºC in Sfp lysis buffer (20 mM Tris-HCl, pH 7.9, 500 mM NaCl, and 10 mM imidazole) supplemented with protease inhibitors The beads were washed with 100 mL of Sfp wash buffer (20 mM Tris-HCl, pH 7.9, 500 mM NaCl, and 30 mM imidazole), and protein was eluted with Sfp elution buffer (20 mM Tris-HCl, pH 7.9, 500 mM NaCl, and 250 mM imidazole). Positive fractions were pooled and dialyzed overnight against Sfp dialysis buffer I (50 mM HEPES-KOH, pH 7.6, 100 mM KCl, and 50 % glycerol). Dialyzed Sfp transferase was then concentrated 12-fold in a 3 kDa Amicon® Ultra-15 Centrifugal Filter Units (Millipore # UFC9003). Aliquots were snap frozen and stored at −80 °C.

##### Protein labelling

***Cdc45$^{LD555}$*:** Cdc45-iS6 was fluorescently labelled by incubating Cdc45-iS6 with Sfp phosphopantetheinyl transferase and LD555-CoA (Lumidyne Technologies, custom synthesis) in a 1:1:5 molar ratio in Cdc45 gel filtration buffer (250 K-phosphate, pH 7.6, 150 mM KOAc, 10 % glycerol, and 0.5 mM DTT) supplemented with 10 mM $MgCl_2$ at RT for 1 h. Cdc45$^{LD555}$ was separated from unincorporated dye and Sfp phosphopantetheinyl transferase by gel filtration in a Superdex 200 Increase 10/300 GL column (Cytiva #15182085). After gel filtration, positive fractions were pooled and concentrated in an Amicon Ultra-4 centrifugal filter Ultracel 30 k (Millipore # UFC803024). Labelling efficiency was measured to be 85 ± 4 % (measured value ± instrumental error) by measuring the absorption at 280 nm and 555 nm.

***dCas9$^{LD555}$*:** S6-dCas9-Halo was fluorescently labelled by incubating S6-dCas9 with Sfp phosphopantetheinyl transferase and LD555-CoA (Lumidyne Technologies, custom synthesis) in a 1:2:10 molar ratio in dCas9 gel filtration buffer (50 mM HEPES/KOH, pH 7.6, 150 mM KCl, 20 % glycerol, and 1 mM DTT) supplemented with 10 mM $MgCl_2$ at RT for 1 h. dCas9$^{LD555}$ was separated from unincorporated dye and Sfp phosphopantetheinyl transferase by gel filtration in a Superdex 200 Increase 10/300 GL column (Cytiva # 15182085).

***Mcm2-7$^{JF646\text{-}Mcm3}$*:** labeling of Mcm2-7$^{Halo\text{-}Mcm3}$ with fluorescent dye JF646 was carried out as previously described[9].

***dCas9$^{JF646}$***: labeling of dCas9-Halo was with fluorescent dye JF646 carried out as previously described[9].

#### Single-molecule Instrumentation and imaging

##### Buffers

***Buffer A*:** 5 mM Tris-HCl pH 7.5, 0.5 mM EDTA, and 1 M NaCl.

***Buffer B*:** 10 mM HEPES-KOH pH 7.6, 1 mM EDTA, and 1 M KOAc.

***Buffer C*:** 10 mM HEPES-KOH pH 7.6, and 1 mM EDTA.

***Loading Buffer*:** 25 mM HEPES-KOH pH 7.6, 100 mM K glutamate, 10 mM MgOAc, 0.02 % NP40 substitute, 10 % glycerol, 2 mM DTT, 100 µg/ml BSA, and 5 mM ATP.

***HSW Buffer***: 25 mM HEPES-KOH pH 7.6, 300 mM KCl, 10 mM MgOAc, 0.02 % NP40 substitute, 10 % glycerol, 1 mM DTT, and 400 µg/ml BSA.

***CMG Buffer***: 25 mM HEPES-KOH pH 7.6, 250 mM K glutamate, 10 mM MgOAc, 0.02 % NP40 substitute, 10 % glycerol, 1 mM DTT, and 400 µg/ml BSA.

***Elution Buffer***: CMG buffer supplemented with 10 mM biotin.

***Imaging Buffers***: CMG buffer supplemented with 2 mM 1,3,5,7 cyclooctatetraene, 2 mM 4-nitrobenzylalchohol, and 2 mM Trolox.

#### DNA functionalization and binding to magnetic beads

20 µg of 23.6 kb plasmid pGL50-ARS1 containing a natural ARS1 origin were linearized overnight with AflII (NEB # R0520L). The resulting 4-nt overhangs TTAA at both ends of the linear DNA were functionalized by incorporating desthiobiotinylated dATP (Jena Bioscience # NU-835-Desthiobio) and digoxigenylated dUTP (Jena Bioscience #NU-803-DIGXL) with Klenow Fragment (3'→5' exo-) (NEB # M0212L); unincorporated nucleotides were removed with Microspin™ S-400 HR spin columns (GE Healthcare # GE27-5140-01). The functionalized DNA was bound overnight at 4 °C to 4 mg of Dynabeads™ M-280 Streptavidin magnetic beads (Invitrogen # 11205D) in Buffer A. After binding, beads were washed twice with Buffer B, twice with Buffer C, and stored at 4 °C in Buffer C. The amount of bound DNA was measured by comparing the concentration of DNA in the supernatant before and after binding, yielding a binding efficiency of 2.3-2.9 mg DNA (~150-190 fmol)/mg beads.

#### Hybrid ensemble and single-molecule assay

##### Ensemble CMG assembly and activation

CMG assembly and activation reactions were carried out in two stages: Mcm2-7 loading and phosphorylation, and CMG assembly and activation. Unless otherwise specified, each step of the reaction was conducted at 30 °C with 800 rpm shaking:

***Mcm2-7 loading and phosphorylation***: 1 mg of magnetic DNA-bound magnetic beads was washed with 200 µl of Loading Buffer, and resuspended in 75 µl of Loading Buffer. To load Mcm2-7 hexamers onto the origin-containing DNA, 35.7 nM ORC, 50 nM Cdc6, and 100 nM Mcm2-7/Cdt1 (or Mcm2-7$^{JF646\text{-}Halo\text{-}Mcm3}$/Cdt1 or Mcm2-7$^{Mcm2(6A)}$/Cdt1) were incubated with the beads for a total of 30 min, but added to the reaction at 0 min, 5 min and 10 min, respectively. Subsequently, 100 nM DDK was added and the reaction incubated for 30 min. The supernatant was then removed, and the beads were washed once with 200 µl of HSW buffer and once with 200 µl of CMG Buffer.

***CMG assembly and activation***: after washing, beads were resuspended in 50 µl of CMG Buffer supplemented with 5 mM ATP. Then, 50 nM Dpb11, 200 nM GINS, 30 nM Polε, 20 nM S-CDK, 50 nM Cdc45$^{LD555}$, 30 nM Sld3/7, 55 nM Sld2, and 10 nM Mcm10 were added to the reaction and incubated for 15 min; For this step, a master mix of all the proteins was made immediately before and incubated on ice. After CMG assembly and activation, the supernatant was removed, and the beads were washed three times with 200 µl of HSW Buffer and once with 100 µl of CMG Buffer. After washing, the assembled DNA-protein complexes were eluted by resuspending the magnetic beads in 200 µl of Elution Buffer, and incubated at RT for 1 h with 800 rpm shaking. The supernatant was then removed and diluted by the addition of 1400 µl of CMG Buffer, and divided into two 700 µl samples for single-molecule imaging.

##### Single-molecule imaging

In general, single-molecule experiments were performed simultaneously on two instruments that combine optical tweezers and confocal microscopy (C-Trap and Q-Trap, LUMICKS); The only exception to this is the two-color colocalization experiments, which were carried out solely in the C-Trap, LUMICKS. Both instruments use a microfluidic chip with five inlets and one outlet. Three of these channels are injected from the left and used for bead trapping and DNA-protein complex-trapping. The other two channels were used as protein reservoirs and buffer exchange locations (Fig. 1a). Prior to each experiment, the microfluidic chips of both instruments were passivated for at least 30 min with 1 mg/mL bovine serum albumin (BSA, NEB # B9000S) followed by 0.5% Pluronic® F-127 (Sigma # P2443).

In all experiments, the channels contained the following solutions:

***Channel 1*****:** 2.06 µm anti-digoxigenin coated polystyrene beads (Spheroteck # DIGP-20-2) diluted 1:50 in PBS.

***Channel 2*****:** CMG-containing DNA eluted from magnetic beads.

***Channel 3*****:** Imaging Buffer

***Channels 4 and 5*****:** Imaging Buffer and Imaging Buffer supplemented with 5 mM ATP or 5 mM ATPγS (Roche # 11162306001).

Prior to each experiment, the trapping laser power was adjusted to achieve a stiffness of 0.3 pN/nm in both traps[6,42]. Then, individual DNA-molecules were trapped between two bead in channel 2, and the tethering of single DNA molecules was confirmed by analysing the force-extension curve[43]. The DNA was then transferred to either channel 4 or channel 5. The distance between both beads was then fixed to achieve a tension of 2 pN, and the DNA was imaged without flow. In all single-color experiments, fluorescent dye LD555 was illuminated with a 561 nm laser at a power of 4 $\mu$W as measured at the objective, and the fluorescence was detected on a single-photon counting detector. 2D confocal scans were performed over an area of 160 x 18 pixels, which covered the entire DNA stretched at a tension of 2 pN and the edges of both beads. Pixel size was set to 50 x 50 nm, illumination time per pixel was set to 0.2 ms, and the frame rate was set to 5 s.

Dual-color experiments were carried out almost identically, with the following differences: 1) fluorescent dyes LD555 and JF646 were simultaneously illuminated with a 561 nm laser at a power of 4 $\mu$W and a 638 nm laser at a power of 12.5 $\mu$W, and 2) the frame rate was set to 0.7 s. The microscopes output HDF5 files that store the confocal scan data, as well as force data and bead location data monitored during the scan.

### Ensemble assays

#### CMG sliding assay

##### DNA template generation

Both 1.4 kb DNA constructs used had one biotinylated end and the same overall sequence containing an ARS1 origin and an *Hpa*II methyltransferase recognition site (CCGG) at the other end. However, only one of the constructs contained a 5-fluoro-2'-deoxy-cytosine within this recognition site to covalently trap the methyltransferase[27,44]. Both constructs were synthesized by PCR using gBlock™ DRM8 (IDT, custom synthesis) as a template and primer pairs DRM_222 and DRM_220 (for the construct without a protein crosslink), or DRM_222 and DRM_218 (for the construct with a protein crosslink). Both reactions were run on a 0.8 % agarose gel, and the appropriate bands were purified from the gel and stored at -20 °C.

##### DNA:protein crosslink formation and binding to magnetic beads

2.5 µg of each DNA construct were incubated at 37 °C overnight with *Hpa*II methyltransferase in a 50:1 protein:DNA molar ratio in CutSmart™ buffer (NEB) supplemented with 10 µM S-adenosylmethionine (NEB #B9003S). Then, each reaction was bound to 1.5 mg of Dynabeads™ M-

280 Streptavidin magnetic beads (Invitrogen #11205D) in Buffer A for 1 h at 37 °C and 1000 rpm shaking. After binding, beads were washed twice with Buffer B, twice with Buffer C, and stored at 4 °C in Buffer C.

##### CMG sliding assay

Unless otherwise specified, every step of the reaction was carried out at 30 °C with 1250 rpm shaking. For each condition, 250 µg of DNA-bound magnetic beads were washed with 50 µl of Loading Buffer. Then, 35.7 nM ORC, 50 nM Cdc6, and 100 nM Mcm2-7/Cdt1 (or Mcm2-7$^{Mcm2(6A)}$/Cdt1) were added and incubated with the DNA-bound beads for 30 min; a master mix of all the proteins was made immediately before addition and incubated on ice. Subsequently, 100 nM DDK was added, and the reaction incubated for 30 min. The supernatant was removed, and the beads were washed once with 50 µl of HSW Buffer 2 (25 mM HEPES-KOH pH 7.6, 500 mM NaCl, 10 mM MgOAc, 0.02 % NP40 substitute, 10 % glycerol, 1 mM DTT, and 400 µg/ml BSA) and once with 50 µl of CMG Buffer. Beads were then resuspended in 50 µl of CMG Buffer supplemented with 5 mM ATP, 50 nM Dpb11, 200 nM GINS, 30 nM Polε, 20 nM S-CDK, 50 nM Cdc45$^{LD555}$, 30 nM Sld3/7, and 55 nM Sld2 and incubated for 5 min; a master mix of all the proteins was made immediately before and incubated on ice. After CMG assembly, the supernatant was removed, and the beads were washed once with 50 µl of HSW Buffer 2 and once with 50 µl of CMG Buffer. After washing, beads were resuspended in 110 µl of HSW Buffer (with 300 mM KCl) with or without 5 mM ATPγS, and incubated at 30 °C with 1250 rpm shaking. At the indicated time points, a 20 µl sample was taken from each reaction, the supernatant removed, and the beads resuspended in 15 µl of MNase Elution Buffer (45 mM HEPES-KOH pH 7.6, 300 mM KOAc, 5 mM MgOAc, 2 mM $CaCl_2$, and 10 % glycerol) supplemented with 0.45 µl of Microccocal nuclease (NEB # M0247S), and incubated at 30 °C for 2 min without shaking. The supernatant was then collected and run on a 4-12 % Bis-Tris polyacrylamide gel. To monitor the amount of bound fluorescent Cdc45, gels were scanned with a green laser on an Amersham Typhoon. Densitometry was performed on ImageJ.

##### Unwinding assay

All ensemble unwinding assays were carried out as previously described[3].

#### Data analysis

##### Overview of data analysis

After taking confocal scans, the resulting raw image data was processed to generate a table containing the spot detections in each frame. These spot detections are connected between frames to produce traces that contain location and intensity information over time.

During the subsequent motion analysis, we fit linear segments to each location trace. The resulting velocities are used to determine whether a trace is static or not. Finally, we determine the type of motion of each non-static trace using anomalous diffusion analysis.

##### Acquiring trace data from raw images

###### Spot detection and tracking

For spot detection we use the scikit-image implementation of a Laplacian of Gaussian (LoG) blob detector[45]. We set the detection radius $r_{LoG}$ to 5 pixels (250 nm); the LoG sigma parameter is then given by $\sigma_{LoG} = r_{LoG} / \sqrt{2}$. The detection threshold is set to 0.5 ADU/pixel$^2$. Detected spots are localized with subpixel resolution by performing Gaussian profile fits on spot intensity projections in both x- and y-directions. For frame-by-frame tracking of the spots, we use our own implementation of the Linear Assignment Problem (LAP) framework[46] with a maximum spot linking distance of 6 pixels (300 nm) and a maximum frame gap of 3 frames (15 s). Spots are considered colocalized if they are less than 2 pixels (100 nm) apart. Spot intensities are given by the total photon count within the detection radius.

#### Location and fluorophore intensity calibration

We use the fluorescent dCas9 data (Extended Data Fig. 3j) to calibrate spot locations and expected fluorophore bleaching step sizes. Because the location of the dCas9 on the DNA is known, a pixel-to-base-pair map can be made for the confocal images by mapping the mean location of all dCas9 spots (on the left and the right side of the DNA) to the corresponding locations in base pairs. Moreover, because dCas9 is labelled with the same fluorophores as the fluorescent proteins used in the CMG experiments, and because dCas9 spots contain one dCas9 molecule, we can find the fluorophore bleaching step size mean $\mu_{\Delta I}$ and standard deviation $\sigma_{\Delta I}$. The minimum bleaching step size, needed for bleaching step fitting, is set to $\Delta I_{min} \leq \mu_{\Delta I}$ - 2 $\sigma_{\Delta I}$ to capture at least 95% of all bleaching events.

#### Determination of number of fluorophores per diffraction-limited spot

To determine the number of fluorescently labelled proteins within each diffraction-limited spot, we count the number of photobleaching step within each spot. For experiments with multiple laser colors (in this case red ($r$) and green ($g$)), we first correct spot intensities for crosstalk by using the equation $I_{r,\ corrected} = I_r - I_g \cdot \mu_{\Delta I,\ r\ (crosstalk)} / \mu_{\Delta I,\ g}$. Then, we fit bleaching traces to a piecewise constant function using Change-Point Analysis (CPA) (we use a Python implementation called 'ruptures'[47]). We use an L2 cost function to detect mean-shifts in the signal, with a minimum segment length of 2 and a penalty term of $\Delta I_{min}^2$. If any steps smaller than $\Delta I_{min}$ are detected, these are pruned starting at the smallest step, until only steps larger than $\Delta I_{min}$ remain.

#### Data filtering

The resulting data table of traces with number of fluorescent proteins per spot was filtered in order to reduce noise, outliers, and data that is not suitable for further motion analysis:

1. While the distance between the optical traps is constant, the force between the traps can fluctuate; jumps in the force signal could indicate, for instance, DNA 'slipping' from the beads, or a protein aggregate landing on a bead, which makes the location signal inaccurate. Hence, if the force signal exhibits a jump larger than $2\sigma_F$ after fitting with CPA, where $\sigma_F = 0.1$ pN is the force fluctuation of a clean trace, only the part of the trace before that jump is used for motion analysis.
2. Diffraction-limited spots containing more than 5 fluorescent proteins, likely aggregates, are filtered out.
3. Any traces starting or ending within 1 kbp from a bead are filtered out to prevent any proteins likely stuck to a bead from entering the dataset.
4. Any traces starting after frame 3 are also filtered away because we do not expect any fluorescent protein to land on the DNA during the scan.
5. The last frame of each trace is omitted for motion analysis because photobleaching often happens while that frame is being taken, resulting in a distorted spot with an incorrect position.
6. Finally, in order to perform reliable motion analysis, only traces with a length of 14 frames or more are retained and used for motion analysis (Extended Data Fig. 6c).

### Positional analysis

In all positional plots, we report the average position of the first three frames of each trace as the initial position of CMG. The bin size of the initial position histograms was set to 700 bp to be close to the diffraction limit while having the ARS1 origin positioned near the center of its corresponding bin.

### Motion analysis

#### Trace segment fitting

To reduce noise before we fit segments to each trace, we first apply a Kalman filter with expectation-maximization (using the pykalman Python library https://github.com/pykalman/pykalman). Then, we fit linear segments with CPA[47], this time using a linear cost function to fit a multiple linear regression model to the trace. The minimum segment size is set to 3 and the penalty term is set to 0.3; halving or doubling the penalty term does not give a significant change in results, showing that the results are robust. After this procedure, each spot detection has associated with it a CPA-fitted velocity $v_{CPA}$.

The CPA fit makes sense for static traces and traces exhibiting piecewise linear motion, but not for spots undergoing diffusive motion. For the latter, we expect to see CPA segments with randomly alternating directions, and random velocities from some distribution with a variance dictated by the diffusion constant. A detailed description of the analysis of diffusive spots can be found below.

#### Motion calibration

The distribution of $v_{CPA}$ for fluorescent dCas9 gives us two values to calibrate the motion analysis. Firstly, the mean velocity $\mu_v$ = 0.38 bp/s provides a drift correction value. Secondly, the standard deviation $\sigma_v$ = 0.40 bp/s gives us a cutoff value to determine whether a diffraction-limited spot is static or not; we set this cut-off at the conservative value of $5\sigma_v$ = 2.0 bp/s.

Another value we need for further analysis is the location measurement error $\sigma_x$. This error is given by the standard deviation of detected dCas9 locations around their mean, after drift correction, which is found to be $\sigma_x$ = 72 bp (≈ 24 nm) (Extended Data Fig. 3e).

#### Anomalous diffusion analysis

First, we correct each trace for drift with $x_{corrected}(t) = x(t) - t \cdot \mu_v$. Then we use mean squared displacement (MSD) analysis[26] to fit an anomalous diffusion exponent $\alpha$, which characterizes the motion type of each mobile trace. The MSD has the form:

$$\mathrm{MSD}(\tau) = D_\alpha \tau^\alpha + 2\,\sigma_x^2 \quad \textbf{(Eq. 1)},$$

where $D_\alpha$ is the anomalous diffusion constant and $\tau$ is the lag time. For spots undergoing confined diffusion, $\alpha \ll 1$; for freely diffusive spots $\alpha \approx 1$, and for traces exhibiting unidirectional motion $\alpha \gg 1$. The fit is performed through the logarithm of the measurement error corrected MSD:

$$\log(\mathrm{MSD}(\tau) - 2\,\sigma_x^2) = \log(D_\alpha) + \alpha \log(\tau) \quad \textbf{(Eq. 2)}.$$

We use least squares to fit up to a maximum lag time $\tau_M$ of 33% of the total length of the trace, with a minimum $\tau_M$ of 5 frames and a maximum of 50 frames. The value of $\alpha$ is constrained between 0 and 2. The trace is then placed into one of three categories using the fitted value of $\alpha$, with confined diffusive spots $0 \leq \alpha < 0.5$, freely diffusive spots $0.5 \leq \alpha < 1.5$, and unidirectionally moving spots $1.5 \leq \alpha \leq 2$. Because we expect populations around $\alpha \approx 1$ and $\alpha \approx 2$, we need the error in alpha $\sigma_\alpha$ to be < 0.5 in order to ensure statistically significant results.

#### Calculation of diffusion coefficients

For traces that are found to be diffusive, we calculate the diffusion coefficient by redoing the MSD fit, setting $\alpha = 1$, and using a previously published appropriate range of delay times[48].

#### Anomalous diffusion exponent error determination

In order to study the error in $\alpha$ as a function of minimum trace length, we have run the same analysis on 512 simulated diffusive traces and 512 simulated traces with a constant speed (with $\alpha = 1$ and $\alpha = 2$, respectively), with representative values for the diffusion constant $D = 1.5 \times 10^{-3}$ kb$^2$/s and speed $v$ = 5 bp/s. We use the experimentally determined measurement error ($\sigma_x$ = 72 bp) and mean fluorophore lifetime (25 frames). These simulations show that we need a minimum trace length of 14 frames for the error in alpha, $\sigma_\alpha$, to be < 0.5, justified by the motion classification cutoffs discussed above.

In all plots we use experimental means and standard deviations whenever possible. On population bar plots we use the statistical error, i.e., the standard error of proportion, given by $\sqrt{p(1-p)/\,n}$, with $p$ the measured proportion and $n$ the sample size.

#### Bin size selections

In general, the bin size of all the histograms in this manuscript were chosen to be larger but in the order of magnitude of the error of the random variable being displayed. Specifically:

- The bin size of the initial position histograms was set to 700 bp to be close to the diffraction limit while having the ARS1 origin positioned near the center of its corresponding bin.
- The bin size of the absolute instantaneous velocities histograms was set to 2.5 bp/s, which is ~ 6 X the velocity noise (Fig. 2a inset)
- The bin size of the absolute mean velocities histograms was set to 1.0 bp/s, which is ~ 2 X the velocity noise (Fig. 2a inset)
- The bin size of instantaneous velocities from CPA histograms was set to 1.0 bp/s, which is ~ 2 X the velocity noise (Fig. 2a inset)
- The bin size of the processivities histograms was set to 0.5 kbp, which is ~ 7 X the location error (Extended Data Fig. 3e)
- The bin sizes of the histograms of step sizes and location errors of dCas9 (Extended Data Fig. 3a-b and e) are irrelevant because we only use the means and the standard deviations of the underlying distribution for our analysis.
- The bin sizes of the histograms of anomalous diffusion exponents $\alpha$ are set to 0.25, which is ~ 1/2 the error in $\alpha$ (Extended Data Fig. 6c). These histograms, however, were only an intermediate in our analysis. In the final analysis, a bin size of 0.5 ~ the error in $\alpha$ (Extended Data Fig. 6c, Methods) was used to classify motion types.

#### Data and code availability

Data can be found at https//doi.org/10.4121/19948253, and contains:

- A table with an overview of experiments.
- Raw tiff images sorted by experimental condition and by setup.
- Filtered spot tracking tables, with connected spot detections for each frame in each scan. Each row has a scan_id and trace_id.
- Motion analysis summary tables, containing motion information for each trace. The scan_id, trace_id fields link this table to the table containing the full trace information.

Code is available at https://gitlab.tudelft.nl/nynke-dekker-lab/private/c-trap/ctrappy and contains:

- Notebooks (at notebooks/notebooks_CMG_paper/) for further analysis on trace tables and for plotting the results.

#### Methods References

#### Tables:

Table 1: oligos and primers used in this study

| Name | 5' to 3' sequence |
|---|---|
| DRM_005 | GCTGCGCCTGCTGAACGGTGATTATAAAGATGATGATGGG |
| DRM_006 | AGCCAGCTCAGGCTATCGCCCTCGTCTGTGACTTCATC |
| DRM_184 | ACGGCTGTTAAATGGGGGGAGTGATAAGAAATACTCAATAGGC |
| DRM_185 | AATAACCAACTTAATGAATCCCCCACGTGATGATGATGATG |
| TL_033 | GCGCGCCAATTGGAGCTCCACCGCGG |
| TL_034 | GGCGCGCCGGAAACAGCTATGACCATGATTACGCC |
| DRM_218 | ATACTTTAGATTGATTTC[5-Fluoro-2'-dC]GGCTTCACCTG |
| DRM_220 | ATACTTTAGATTGATTTCCGGCTTCACCTG |
| DRM_222 | Biotin-CTAGTGGATCCCCAGGGCT |
